# Multi-modal atlas of lifestyle interventions reveals malleability of ageing-linked molecular features

**DOI:** 10.1101/2025.08.30.673115

**Authors:** Chiara MS Herzog, Charlotte Vavourakis, Elisa Redl, Magdalena Hagen, Gabriel Knoll, Christina Watschinger, Umesh Kumar, Bente Theeuwes, Juliane Gasser, Sophia Zollner-Kiechl, Robert Reihs, Heimo Müller, Maria Cavinato, Sonja Sturm, Hermann Stuppner, Alexander Moschen, Birgit Weinberger, Tobias Greitemeyer, Matthias Schmuth, Verena Moosbrugger-Martinz, Thomas Trafoier, Verena Lindner, Anna Wimmer, Peter Widschwendter, Hans-Peter Platzer, Alexander Höller, Michael Knoflach, Wolfgang Schobersberger, Martin Widschwendter

**Author notes:** Contact information: Chiara Herzog and Martin Widschwendter.

## Abstract

Extending human healthspan requires understanding how lifestyle interventions impact molecular systems across tissues and time. Here, we present the TirolGESUND Lifestyle Atlas (ClinicalTrials.gov: NCT05678426), a longitudinal, multi-modal resource profiling 156 healthy women (aged 30-60 years) undergoing 6-month intermittent fasting (n=114) or smoking cessation (n=42) interventions. Participants were sampled up to four times across seven tissues and fluids, generating >3,450 biospecimens with harmonised DNA methylation, metabolomics, microbiome, and immune profiling, alongside skin histology, barrier measurements, and rich clinical metadata. We demonstrate the utility of this dataset through: (i) multi-omics-wide association studies linking traits to molecular features; (ii) integrative factor modelling revealing coordinated cross-tissue signatures; (iii) epigenetic-biomarker cross-omic associations, and (iv) CpG-level variance decomposition mapping stable, individual-specific, tissue-restricted, and intervention-responsive methylation patterns. We further show that ageing-linked features are selectively malleable: highly compliant intermittent fasting participants exhibited attenuated or even age-opposing molecular trajectories within six months. The atlas enables unprecedented within-cohort comparisons across omic layers and tissues, supporting discovery of context-dependent biomarkers, cross-system coordination, and intervention responsiveness. Data are available via an interactive portal, with sensitive data under controlled access (https://eutops.github.io/lifestyle-atlas/). This resource provides a foundation for exploring biomarker association and multi-tissue epigenetics, enabling hypothesis generation and benchmarking for systems biology and human healthspan research.

**Graphical abstract:** 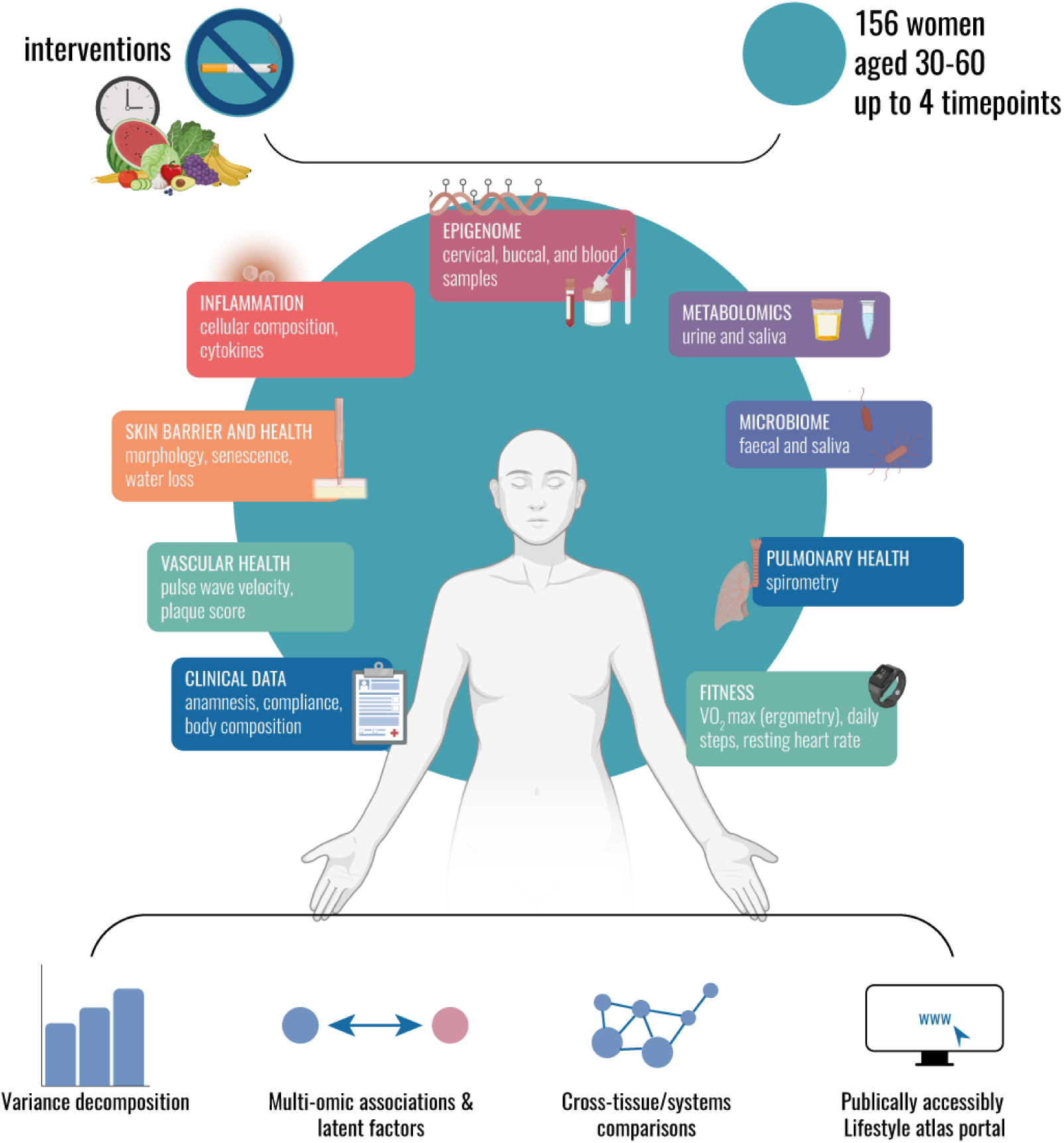

**Highlights:**

- Multi-tissue multi-modal profiling of two clinically-relevant lifestyle interventions in 156 women aged 30-60
- Intermittent fasting modifies ageing-linked molecular features in an age-opposing direction
- Cross-tissue epigenetic biomarker mapping links immunity, metabolism, and microbiome
- Tissue-specific epigenetic plasticity maps reveal candidate meQTLs,
- Data sharing and interactive portal enable broad reuse for hypothesis generation and exploration

## Introduction

Extending healthspan - defined as the period prior to onset of chronic diseases and disabilities of ageing ^1^ - is an urgent priority in the context of global population ageing. Lifestyle factors such as poor diet, obesity, and tobacco smoking are well-known contributors to disease risk ^2,3^ and have been implicated in accelerating cellular ageing processes ^4–7^. While lifestyle interventions like dietary restriction and smoking cessation offer promising, accessible strategies to reduce long-term disease risk and extend healthspan, our ability to assess their systemic biological impact in humans remains limited by the lack of robust surrogate biomarkers and comprehensive longitudinal datasets spanning multiple tissues and molecular systems.

Recent advances in molecular profiling have yielded promising candidate biomarkers, including DNA methylation (DNAme)-based epigenetic clocks ^8–12^, cancer risk biomarkers ^13–16^, and exposure metrics ^17^, as well as microbiome, metabolic, and immune signatures, to monitor various aspects of health. However, most human studies capture only one dimension of a complex picture: multi-omic profiling to a single tissue at a single time point, or longitudinal profiling of a single omic layer, often without paired clinical or intervention metadata. While such datasets have advanced biomarker discovery, they offer only partial insight into the systemic biological changes that accompany healthspan-extending interventions. In particular, no existing resource integrates multiple omic layers, tissue compartments, and timepoints in the context of clinically relevant lifestyle perturbations, limiting our ability to compare system-specific responsiveness, identify cross-system molecular coordination, or determine which tissues most sensitively reflect meaningful health changes. Previous efforts such as iPOP ^18,19^, Arivale ^20^, CALERIE ^21^, the Human Phenotype Project ^22^, a Finnish study by Marabita et al. ^23^, and recent work exploring the impact of exercise by Geng et al. ^24^, illustrate the value of longitudinal or multi-omic profiling, but to our knowledge none integrate all these dimensions (time, multi-tissue (>3), multi-omic, multiple lifestyle interventions) alongside rich clinical phenotyping.

To address this gap, we developed a multi-modal longitudinal lifestyle atlas, a publicly accessible resource generated from the TirolGESUND study (ClinicalTrials.gov: NCT05678426). The study enrolled 156 healthy women aged 30-60 years into two parallel, clinically relevant 6-month interventions - intermittent fasting (n=114) and smoking cessation (n=42) - and profiled participants longitudinally using multi-tissue sampling (seven tissue/fluid types) across up to four timepoints per participant, generating over 3,450 biospecimens linked to harmonised multi-omic, clinical, and lifestyle metadata. Biospecimens include buccal, cervical, blood, saliva, faecal, urine, and skin tissue, enabling comprehensive analysis of the epigenome, microbiome, metabolome, and immune system across time and tissues. Extensive clinical, epidemiological, and lifestyle metadata support stratification by age, body-mass index, menopause status, and intervention response. A set of skin biopsies profiled using histology further supports analysis of how accessible molecular biomarkers reflect deeper tissue states.

Here, we introduce a longitudinal study framework specifically designed to enable integrated, cross-omic analyses over time, required for system-level exploration of health. Illustrative applications include multi-omic-wide association studies (MOWAS) linking clinical traits to molecular features, integrative factor modelling to uncover coordinated sources of variation at baseline and longitudinally, cross-omic correlation analyses of epigenetic biomarkers, and CpG-level variance decomposition to map stable, individual-, tissue-, and intervention-specific methylation patterns. Designed from the outset as a reusable community resource for biomarker discovery, systems biology, and method benchmarking, the dataset is available in accordance with FAIR principles via an interactive data portal. We anticipate broad reuse of this resource across systems biology and biomarker development and validation, and envision it as a foundational model for future multi-modal, multi-omic studies of healthspan and disease prevention.

## Results

### A longitudinal multi-tissue, multi-omic resource for systemic and system-specific analysis of lifestyle interventions

To investigate how systemic molecular profiles respond to healthspan-promoting interventions in a generally healthy but at-risk population, we established a longitudinal, multi-omic intervention study in which participants underwent one of two real-world behavioural interventions. The TirolGESUND study enrolled 156 women aged 30 to 60 years into one of two parallel six-month interventions based on lifestyle risk factors: intermittent fasting (n=114, BMI ≥25) or smoking cessation (n=42, ≥10 cigarettes/day for ≥5 years) (**Figure 1a**). These interventions target distinct but complementary biological pathways (energy metabolism and nutrient signalling [intermittent fasting] versus toxin exposure [smoking cessation]), enabling comparative analyses of systemic responses.

**Figure 1.**
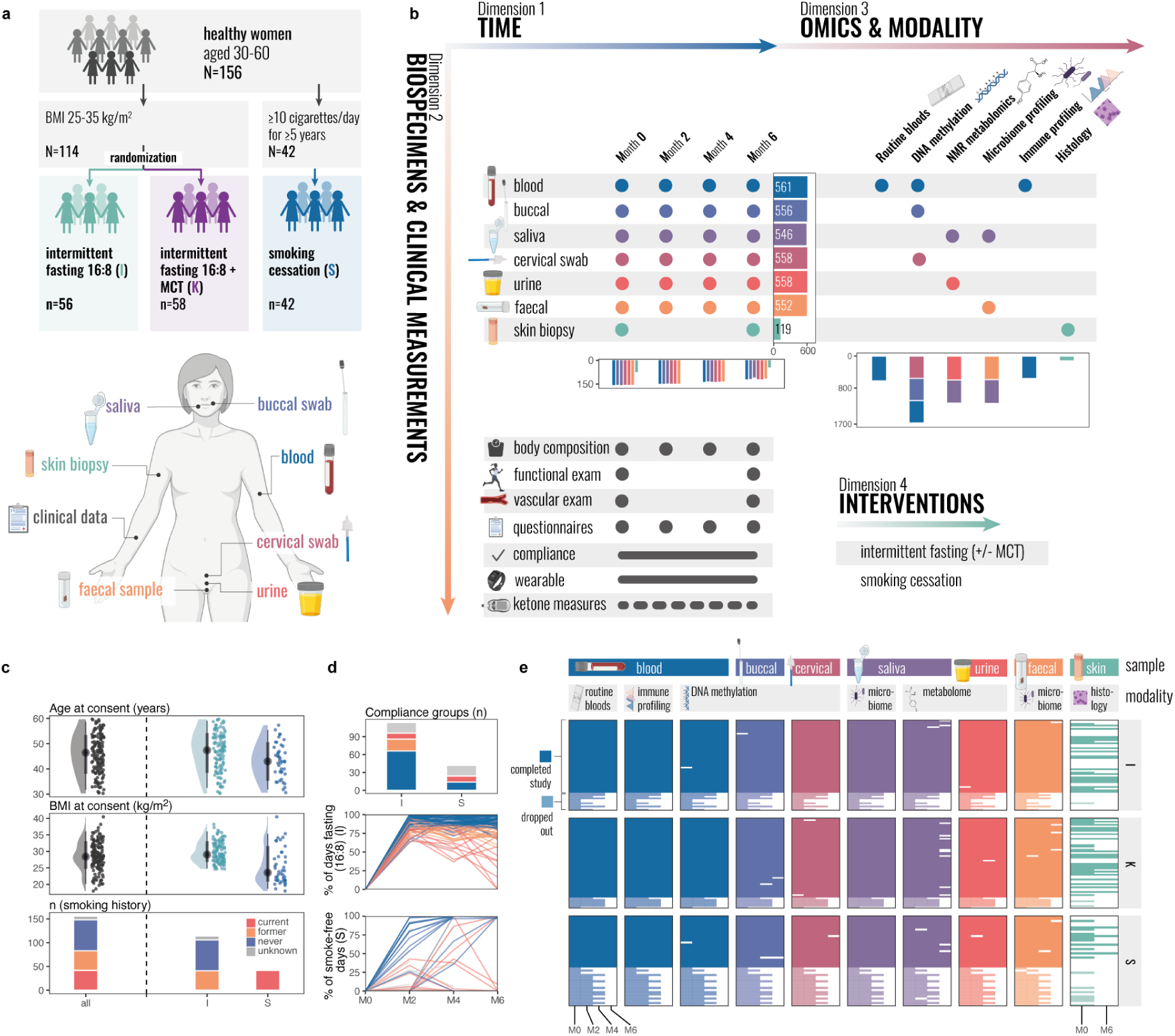
Overview of the TirolGESUND study design and dataset structure. **a** Participants were enrolled into two parallel lifestyle intervention arms, intermittent fasting with or without ketogenic supplement (medium chain triglycerides, MCT) (n=114) and smoking cessation (n=42). **b** Biospecimens from seven tissue and fluid types were collected at each timepoint and profiled using multiple omics technologies, enabling multi-layer, multi-tissue longitudinal analyses. The resulting dataset spans four key dimensions: (1) time - participants were followed for 6 months with four study visits; (2) biospecimens and clinical measurements - multiple sample types were collected at each visit (total numbers per sample type and visit are visualised in the bottom panel panel); (3) omics and phenotyping modalities - each specimen was profiled using one or more molecular assays (total number per assay shown in bottom panel); and (4) interventions - two lifestyle intervention arms acting as biological perturbations. **c** Baseline characteristics of all participants and stratified by study arm (“I”, dietary intervention; “S”, smoking cessation), including age, body-mass index, and smoking history. **d** Participant compliance profiles, stratified by intervention arm and grouped by longitudinal adherence level (bottom two panels). The intervention was initiated immediately after M0. **e** Data availability matrix per participant (rows) and sample type/modality (columns), stratified by intervention. White fields indicate missing samples. Transparent boxes denote participation dropout. **Abbreviations**: MCT, medium-chain triglycerides. I, intermittent fasting study arm. S, smoking cessation study arm. M0-M6, month 0-6.

Participants were followed across four study visits (baseline, month 2, 4, and 6), with optional follow-up visits at month 12 and 18 (**Figure S1**). Biospecimens from seven tissue and fluid types - blood, buccal swabs, cervical swabs, saliva, urine, and stool, and, in a subset, skin biopsies - were collected longitudinally and profiled using a suite of molecular technologies capturing multiple biological layers, including DNA methylation, microbiome composition, metabolomics, quantitative and functional immune cell profiling (abundance and cytokine activation following stimulation), and tissue histology. This design enables both compartment-specific analyses (e.g., within the immune system) and cross-layer, cross-tissue comparisons, capturing coordinated or distinct molecular responses to lifestyle interventions. In parallel, structured clinical assessments, questionnaires, and wearable/digital data generated a multi-dimensional dataset integrating molecular, physiological, and lifestyle information across time, tissue, and study arm (**Figure 1b**). Baseline participant demographics are shown in **Figure 1c**.

The study employed a baseline-controlled longitudinal design, focusing on within-individual changes. Participants were not excluded for non-compliance. Instead, adherence was continuously monitored and quantified throughout the study, allowing for dose-response analyses and nested comparisons between compliant and non-compliant individuals (**Figure 1d**). Overall, participant engagement was high, with many individuals, particularly in the dietary intervention arm, demonstrating sustained adherence over the full 6-month period.

In total, we collected 3,450 biological samples and >500 participant-visits, generating a high-density dataset of matched clinical, behaviour, wearable/digital, and multi-omic molecular profiles (**Figure 1b, e**). All data were harmonised, technically validated, and quality-checked across timepoints (**Figure S2-S4**). Data availability across sample types and omic layers is shown in **Figure 1e** and an overview of data hierarchy and structure is provided in **Figure S5**. Data and findings are available through our data portal (see Data and Code Availability, https://eutops.github.io/lifestyle-atlas/), enabling broad reuse across research, biomarker exploration, and systems medicine.

### Different omics capture stable versus dynamic aspects of individual biology

To characterise the behaviour of features in the dataset, we performed variance decomposition using linear mixed models across major omic modalities, clinical variables, and select methylation biomarkers (see **Table S1** for the full list of variables and biomarkers included). For each feature, the total variance was partitioned into components attributable to study arm, inter-individual differences, and intra-individual longitudinal variation (including unmeasured sources of temporal variability). Despite differences in BMI and smoking history between the groups allocated to either dietary or smoking cessation interventions (**Figure 1c**), the study arm (dietary or smoking cessation) explained only a minor proportion of variance (typically <10%; **Figure 2a**, **Table S2**), consistent with much of the variance being captured by baseline inter-individual differences between arms, which are further explored in later analyses. The dominant sources of variation were inter- and intra-individual differences, but their relative contributions varied markedly across modalities. Metabolomic and microbiome features (both at the amplicon sequence variant (ASV)- and family level) exhibited high intra-individual variability, consistent with their sensitivity to short-term exposures, such as recent dietary intake. In contrast, clinical phenotypes and immune cell composition were more personally stable, with inter-individual variation accounting for the majority of variance in these features. DNA methylation biomarkers, including 55 composite scores quantifying ageing, smoking, mortality, or cancer risk using blood, cervical, or buccal methylation data (e.g., GrimAgeAccel (V2) ^25^, a blood biomarker associated with mortality risk; the WID-buccal-BC ^16^, a buccal biomarker associated with breast cancer risk; or the WID-OC, a cervical biomarker associated with ovarian cancer risk ^14^), showed a balanced contribution of intra- and interindividual variance, exhibiting roughly equal variability across individuals and time, suggesting that these features capture both stable traits and responsive molecular states (**Figure 2a**). Cervical and buccal methylation biomarkers exhibited lower inter-individual variability than blood methylation biomarkers. We also explored the variance distribution across the individual CpG sites on the Illumina HumanMethylationEPIC array (version 1), focusing on the most reliable 10% of probes, leveraging a previously described method ^26^ that applies probe-level simulation of the influence of technical noise on methylation β values using the background intensities of negative control probes (see STAR Methods for details). This showed that these sites exhibited similar distributions across inter- and intra-individual distributions (**Figure 2b**).

**Figure 2.**
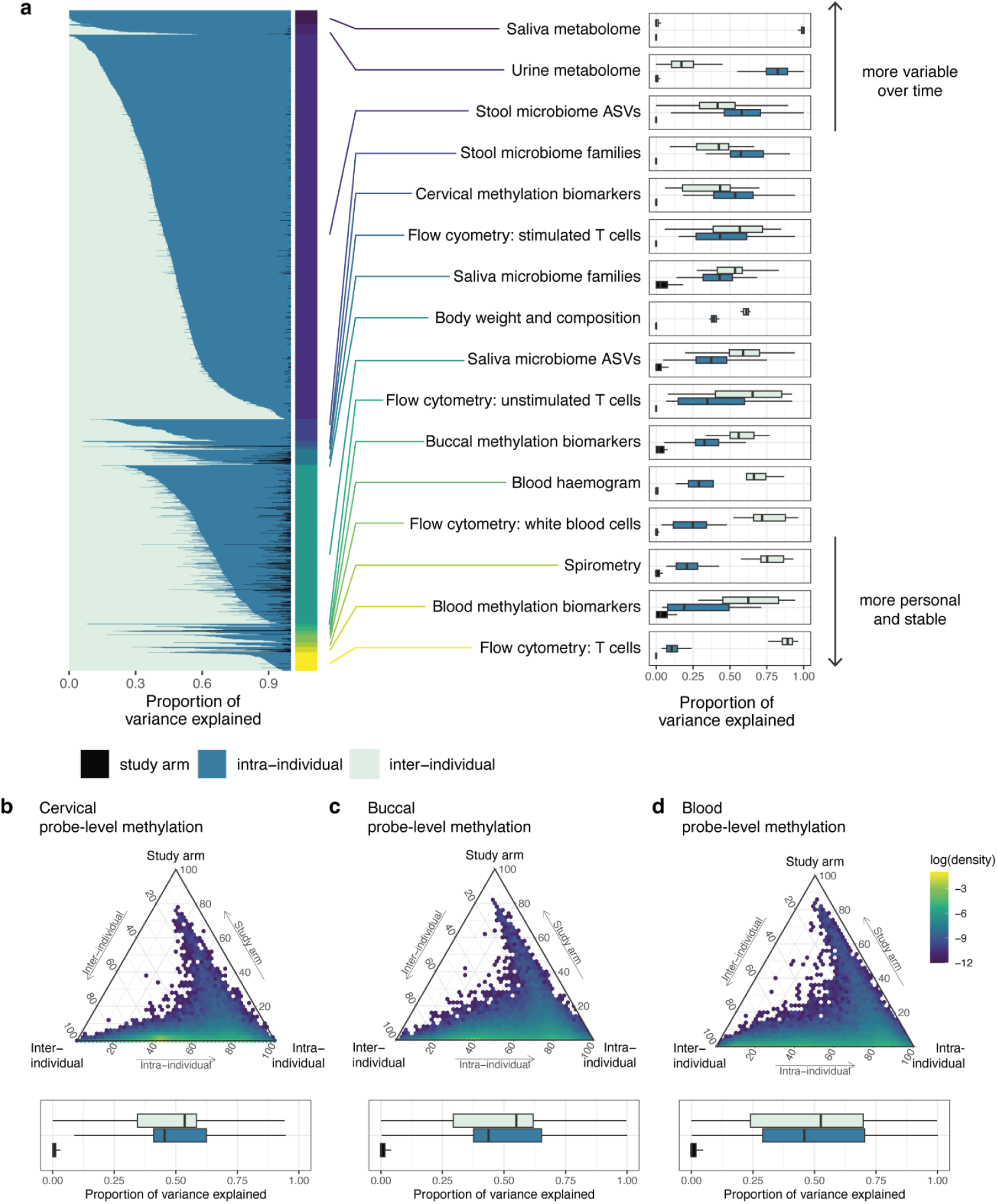
Variance decomposition of omic and clinical data layers highlights distinct temporal variability. **a** Variance was decomposed into study arm, inter-individual variability, and intra-individual (residual) variability using linear mixed models. **b** Probe-level variance decomposition of methylation data on the Illumina HumanMethylationEPIC array (version 1). The top 10% reliable probes, defined using a dynamic thresholding method estimating the impact of technical noise on methylation β values, were analysed (see STAR Methods). **Abbreviations**: ASV, amplicon sequence variant.

These findings reflect fundamental differences in biological variability across molecular systems and underscore the value of measuring multiple omic layers to monitor both stable and dynamic aspects of individual biology.

### Specific modalities preferentially capture distinct population traits at baseline

We next assessed how well molecular features captured inter-individual differences in key phenotypes at baseline, performing a multi-omic wide association study (MOWAS) across all features to explore associations with chronological age, menopausal status, smoking behaviour (current and ever), BMI, VO_2_peak, and alcohol consumption (weekly units at baseline). Distinct patterns of trait associations emerged across modalities (**Figure 3a-g**). Chronological age and smoking behaviours (current, ever) exhibited the highest number of significant associations across features (n=2,716, n=6590, and n=1,002, respectively). As expected, chronological age was captured across multiple layers and reflected strongly across DNA methylation (both individual features and composite biomarkers), immune cell composition (flow cytometry), and clinical measurements (**Figure 3a**), with proportionally fewer associations in metabolomic and microbiome features - consistent with their more dynamic, exposure-responsive profiles (**Figure 2a**). Both current and ever smoking were associated with changes in salivary microbiome composition and, as previously reported, methylation levels (**Figure 3b, c**). VO_2_peak, BMI, alcohol use, and menopause exhibited fewer associations (**Figure 3d-g**).

**Figure 3.**
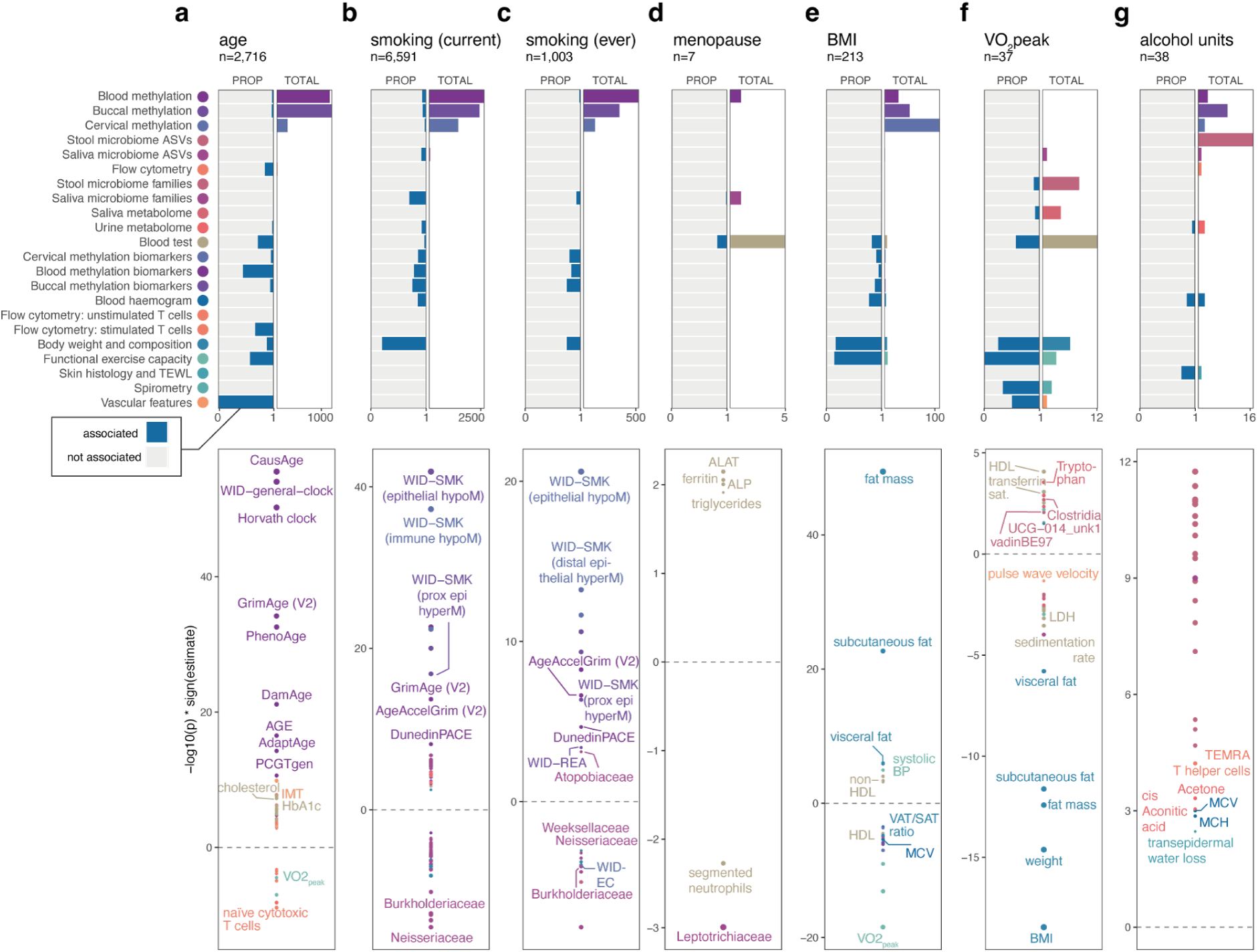
Multi-omic-wide association study (MOWAS) analysis at baseline. Proportion out of total features that were significant (PROP), total number of significant features (TOTAL), and top associations per layer associated with **a** chronological age, **b** current smoking, **c** ever smoking, **d** menopause, **e** BMI, **f** VO2_peak_, and **g** alcohol units. Features significantly associated with the variable at FDR<0.05 (FDR<0.1 for menopause and VO_2_peak) within each layer are shown. Visualisation of associations (bottom panels) exclude individual CpG sites and ASVs for clarity. Full results, including individual CpG sites and ASVs, are presented in **Table S3** and on the interactive portal. **Abbreviations**. FDR, false discovery rate. PROP, proportion of total features significant. BMI, body mass index. ASV, amplicon sequence variant. TEWL, transepidermal water loss. IMT, intima-media thickness. LDH, lactate dehydrogenase. MCV, mean cellular volume. MCH, mean cellular haemoglobin.

To explore these associations in greater detail, we identified the top-ranking features for each trait and visualised them, excluding individual CpG sites and ASVs for clarity, revealing a combination of well-established and novel trait-feature relationships across molecular systems. For instance, age was positively associated with vascular and metabolic parameters (**Figure 3a**), including intima-media thickness (IMT), HbA1c, and cholesterol and negatively associated with VO_2_peak and naive cytotoxic T cells. As expected, several epigenetic biomarkers were strongly associated with chronological age, including CausAge, the WID-general-clock, the Horvath clock, GrimAge V2, and others. Notably, AgeAccelGrim exhibited a negative association with chronological age, in our study population. Both current and ever smoking were strongly associated with smoking biomarkers (**Figure 3b, c**), but also with AgeAccelGrim, DunedinPACE, and several salivary microbiome families (*Burkholderiaceae*, *Neisseriaceae*, *Atopobiaceae*). Menopause was associated with elevated ALAT and ALP levels - previously linked to bone turnover ^27^ - and higher ferritin levels, consistent with cessation of menstruation (**Figure 3d**). Interestingly, a negative association with salivary *Leptotrichiaceae* was also observed. While little is known about the oral microbiome and menopause, these microbes have reported to be sensitive to shifts in estradiol ^28^. BMI was strongly reflected in clinical parameters, including body composition measures, blood pressure, mean cellular volume, and visceral/subcutaneous adipose tissue ratios (VAT/SAT ratio), and, although to a proportionally lesser extent, methylation loci (**Figure 3e**). VO2_peak_ was associated with both clinical measures (HDL, transferrin saturation, LDH, sedimentation rate, pulse wave velocity, and others), microbial taxa, and metabolites, including salivary tryptophan (**Figure 3f**). VO2_peak_ also exhibited a positive association with faecal microbiota *Clostridia UCG-014* and *vadinBE97*. Alcohol consumption was primarily associated with several faecal microbiome strains (ASVs), but also exhibited positive associations with TEMRA T_H_ cells, MCH and MCV, urine cis-aconitate and acetone, and transepidermal water loss, suggesting higher alcohol consumption may lead to decreased skin barrier function.

These results highlight the multi-compartmental nature of lifestyle and ageing-related traits, and demonstrate how different omic layers and modalities contribute complementary insights into human health variation. The full list of significant results is provided in **Table S3** and on our interactive data exploration portal (https://eutops.github.io/lifestyle-atlas/docs/explore/mowas.html).

### Integrative multi-omic factor analysis identifies smoking and age as dominant sources of baseline molecular variation

While feature-level analyses provide insights into associations with phenotypes and interventions, they do not comprehensively capture coordinated patterns across molecular layers. To uncover higher-order structure and shared sources of variation across layers, we applied multi-omic factor analysis (MOFA) ^29^. We first trained MOFA on baseline samples to capture covariation structure independent of longitudinal dynamics. As expected, smoking status and chronological age emerged as the primary axes of molecular variation at baseline (**Figure 4a-c**, **Table S4**), echoing the key sources of variation identified in the baseline MOWAS (**Figure 3**) and consistent with extensive prior literature. Their recovery across tissues and omic layers served as an important internal control, validating the capacity of MOFA to capture biologically interpretable signals across omics. Factor 1 was driven by smoking-associated methylation biomarkers across buccal, blood, and cervical tissues, alongside saliva microbiome families (**Figure 4a, c, Table S5**). Participants in the smoking arm showed a wider spread in Factor 1 than those in the intermittent fasting arm, suggesting it reflects smoking intensity. This was supported by its association with cigarettes per day (**Figure S6a**), although it also correlated with BMI, which was more pronounced in the smoking study arm than in the intermittent fasting arm (**Figure S6b**). Factor 2 was primarily driven by epigenetic ageing markers in blood and immune cells, for instance reduced naive T cells and increased exhausted T cells, consistent with known age-related immune remodelling. It was highly correlated with chronological age (**Figure S6c**) and with intima-media thickness, an established age-related trait (**Figure S6d**). These first two latent factors thus reflect canonical, biologically grounded variation providing a foundation for interpreting less characterised sources of variance explored further below. All weights and associations are provided in **Tables S4 and S5**.

**Figure 4.**
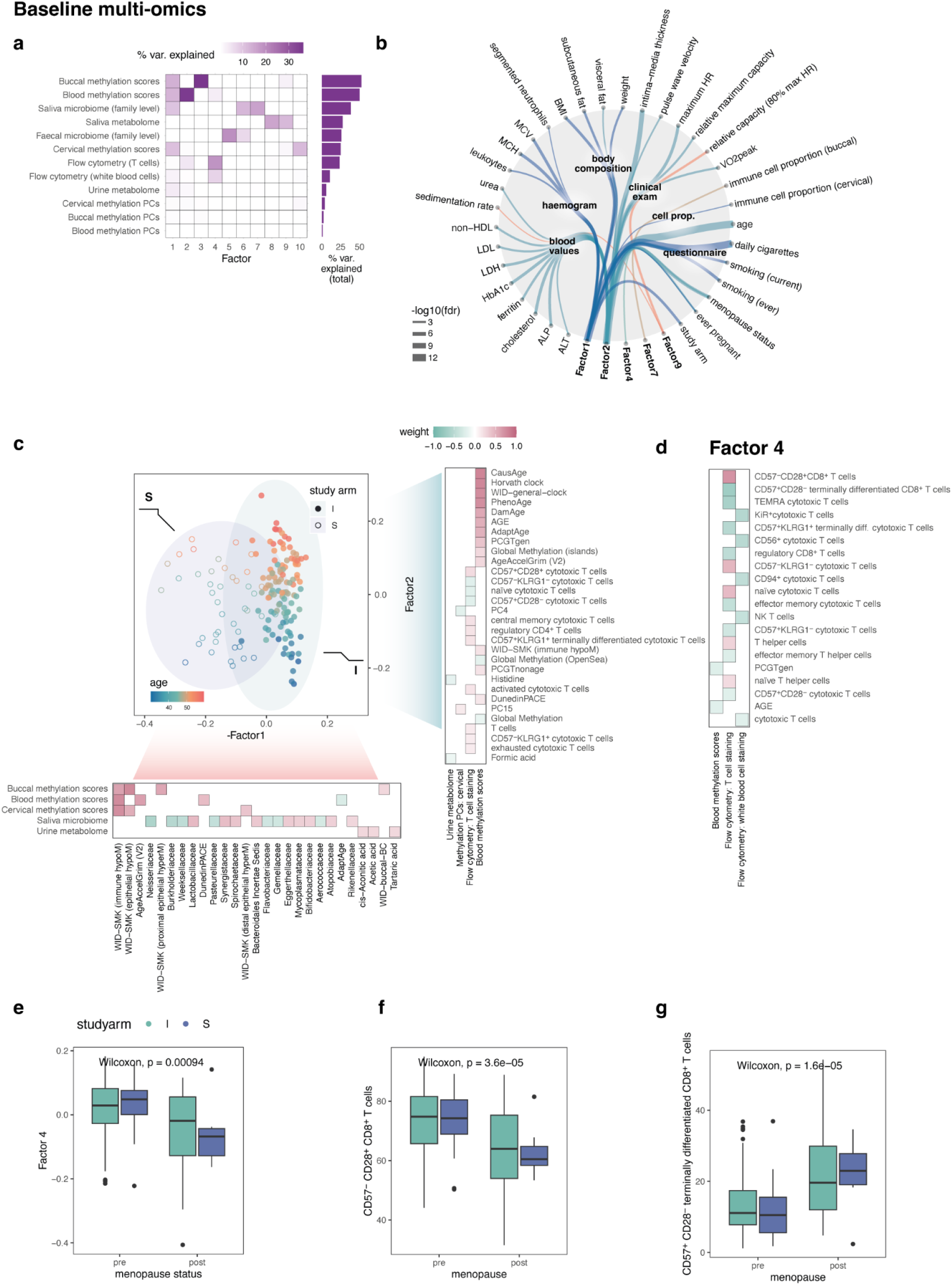
Integrative multi-omic factor analysis (MOFA) at baseline identifies latent factors capturing shared variation across omic layers. **a** Variance explained for each ome by latent factor (heatmap) and overall (side panel). **b** Network diagram of significant associations of baseline multi-omic latent factors with additional recorded phenotypic variables at FDR<0.1. Factors without any significant associations are omitted from the diagram. **c** Scatterplot of individuals by factor 1 and factor 2 at baseline, coloured by age at consent and study arm (intermittent fasting or smoking cessation). Side panels indicate top 30 weights for each factor. **d** Top weights for factor 4, significantly associated with menopause. **e** Boxplot of Factor 4 by menopause status and intervention group. p value is derived from overall Wilcoxon test (pre versus post). **f** Boxplot of CD57^-^ CD28^+^ CD8^+^ T cells, the top contributor to Factor 4, by menopause status and intervention group. p value is derived from overall Wilcoxon test (pre versus post). **g** Boxplot of CD57^+^ CD28^-^ terminally differentiated CD8^+^ T cells, the second-biggest contributor to Factor 4, by menopause status and intervention group. p value is derived from overall Wilcoxon test (pre versus post). **Abbreviations**: PCs, principal component. prop., proportion. ALT, alanine transaminase. ALP, alkaline phosphatase. CRP, c-reactive protein. LDH, lactate dehydrogenase. LDL, low-density lipoprotein. HDL, high-density lipoprotein. MCH, major cellular haemoglobin. MCV, mean corpuscular volume. BMI, body-mass index. HR, heart rate. FVC, forced vital capacity. FEV1, forced expiratory volume after 1 second.

### Menopause associates with immune cell remodelling independent of chronological age

Factor 4 was significantly associated with menopause, but not chronological age, suggesting it reflects menopause-specific changes (**Figure 4b, d, e**). Top weights included immune subsets (**Table S5**): CD57^-^ CD28^+^ CD8^+^ T cells positively associated with Factor 4 (**Figure 4d**, **Figure S7a**) and were reduced in postmenopausal women (**Figure 4f**, **Figure S7b**), while CD57^+^ CD28^-^ terminally differentiated CD8^+^ T cells were negatively associated (**Figure 4d**, **S7c**) and enriched after menopause (**Figure 4g**, **S7d**). Although some age association was observed (**Figure S7c**, **d**), these shifts were more pronounced around menopause (average age 52), rather than following a strictly linear trajectory. Heteroskedasticity in CD57^+^ CD28^-^ terminally differentiated CD8^+^ T cells further supports a menopause-related inflection point in immune remodelling.

### Oral microbiota and metabolites track inflammation and epithelial immune infiltration

Factor 7 negatively correlated with estimated immune proportion in buccal cells, and was dominated by saliva microbiome families (**Figure 4b**, **Figure S8a, b**, **Table S5**), suggesting an interplay between immune presence and microbial composition. Factor 9, composed of saliva metabolites, associated negatively with erythrocyte sedimentation rate (ESR), a systemic inflammation marker, and positively with exercise capacity (**Figure S8c-e**). These findings are consistent with literature linking physical activity to lower ESR ^30,31^ (**Figure S8c, d**). Top contributors were enriched for metabolic pathways including branched-chain amino acid (BCAA) valine, leucine and isoleucine biosynthesis, glycine, serine, and threonine metabolism, and the one carbon pool by folate metabolism (**Figure S8f**), suggesting these metabolite profiles may reflect fitness- or diet-related molecular states. The lack of overlap with microbial families in Factor 9 indicates that these metabolite changes likely arise from host processes rather than microbiota. The ability to detect these patterns was uniquely enabled by the simultaneous profiling of multiple layers in the same individuals.

### Chronological age, menopause, and smoking behaviour shape major axes of molecular change over six months

We next explored latent molecular trajectories over time, applying the longitudinal extension of multi-omic factor analysis (MEFISTO ^32^) across participants and omics layers to explore shared, coordinated trajectories over the course of the study. While several factors integrated signals across omics (e.g., Factors 1, 2, 4, 6), others were dominated driven by individual omic layers, such as blood or buccal methylation (e.g., Factors 3 or 11), or the saliva microbiome (Factor 10) (**Figure 5a, Table S6**). Low inter-factor correlations (**Figure 5b**) indicated these latent factors represented distinct biological axes. Similar to baseline, MEFISTO recovered known ageing and smoking-related features, providing internal validation of model fidelity across time and omic layers. Of note, Factor 2 captured smoking behaviour (**Figure 5c, d**, **Table S7**), but was also responsive to recent smoking cessation (**Figure 5e**), aligning with our findings on the partial molecular reversibility of smoking exposure ^33^. Likewise, Factor 22, driven primarily by the faecal and oral microbiome, was correlated with daily cigarettes in smokers (**Figure S9a-c**).

**Figure 5.**
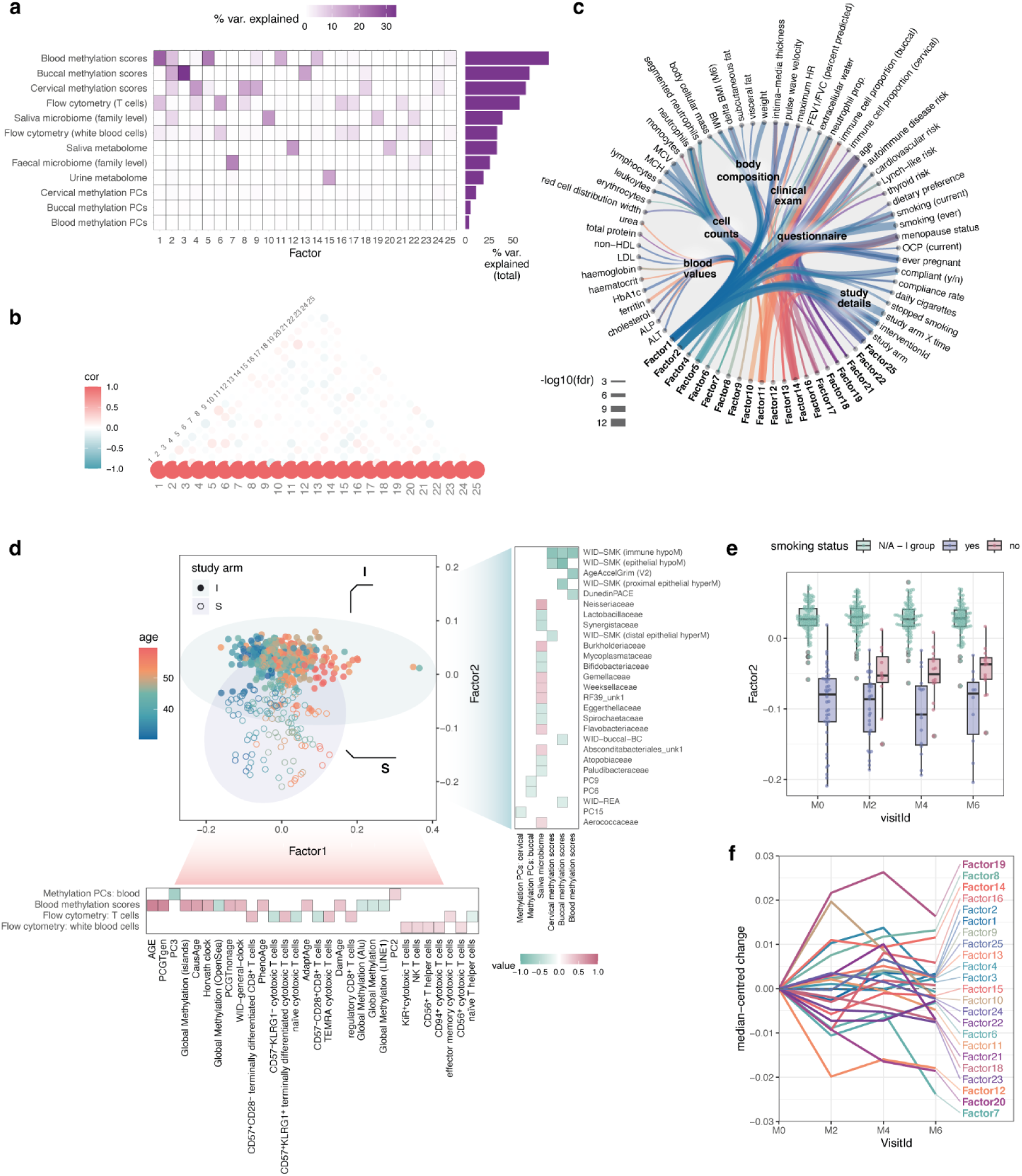
Longitudinal multi-omic factor analysis (MEFISTO) identifies latent factors capturing shared variation across omes over time. **a** Variance explained for each ome by latent factor (heatmap) and overall (side panel). **b** Correlation across factor values. **c** Network diagram of significant associations of multi-omic factors with phenotypic variables at FDR<0.05. Factors without any significant associations are omitted from the diagram. **d** Scatterplot of individuals by factor 1 and factor 2, coloured by age at consent and study arm (intermittent fasting or smoking cessation). Side panels indicate top 30 weights for each factor. **e** Factor 2 values by current smoking status over time in the smoking group (no, yes) and intermittent fasting (I) groups (grouped separately and labelled N/A). **f** Median-centred change in Factor values by visitId. **Abbreviations**: PCs, principal component. prop., proportion. cor, correlation. fdr, false discovery rate. ALT, alanine transaminase. ALP, alkaline phosphatase. CRP, c-reactive protein. HDL, high-density lipoprotein. LDH, lactate dehydrogenase. LDL, low-density lipoprotein. MCH, major cellular haemoglobin. MCV, mean corpuscular volume. BMI, body-mass index. delta BMI (M6), change in body-mass index at month 6 relative to baseline. HR, heart rate. FVC, forced vital capacity. FEV1, forced expiratory volume after 1 second. OCP, oral contraceptive pill use. IF, intermittent fasting.

Factor 5 - capturing a range of blood and cervical age-related biomarkers as well as monocyte populations - exhibited an intriguing association with menopause, skin barrier and transepidermal water loss, segmented neutrophil and total neutrophil counts, diet (**Table S7**, **Figure S9d-h**). Non-classical monocytes were positively associated with Factor 5 and increasing with age (**Figure S9g, i**), in line with recent reports of elevated non-classical monocyte counts in older females ^34^.

These findings reinforced smoking, age and menopause as dominant contributors to molecular variability, both at baseline and over time. Intriguingly, although factors exhibited low correlation, ageing was significantly associated with multiple factors (**Figure 5c**, **Table S7**; Factors 1, 10, 11, 14, 16, 21, and 25), suggesting it exerts widespread but multifaceted effects across molecular layers across longitudinal samples.

### Longitudinal multi-omic shifts over 6 months reflect mixed ageing and intervention-driven influences

We next examined whether any latent factors showed coordinated temporal trends across the entire study population, irrespective of intervention. As expected, most factors remained largely stable over time (**Figure 5f**), consistent with the relative stability of many molecular features (**Figure 2a**) and with the likelihood that distinct interventions elicit divergent trajectories rather than uniform changes. This was also supported by the observation that some alterations were only evident in subgroups of participants (e.g., **Figure 5e**, **Figure S9c**), highlighting the need for content-aware analysis, presented in accompanying manuscripts ^33,35^. Nevertheless, a subset of factors showed overall longitudinal trends. Factors 19, 8, and 14 increased over time, while Factors 12, 20, and 7 declined (**Figure 5f**).

Factor 8 combined methylation and immune parameters (**Figure S10a**, **Table S6**). Its loadings opposed baseline associations with chronological age, consistent with a modest age-opposing effect. Its top association were neutrophil counts and ever smoking status, consistent with a predominantly subgroup-driven effect (e.g., global island methylation increased particularly with smoking cessation, [**Figure S10b**], while CD57^+^CD28^+^ cytotoxic T cells decreased more strongly in the same group [**Figure S10c**]). Factor 19, dominated by oral microbiome features, contained weights both concordant and discordant with chronological age (**Figure S10d**). Its strongest association was buccal immune cell composition (**Table S6**), which shifted most markedly in the intermittent fasting arm (**Figure S10e**). Factor 14 captured several blood methylation biomarkers (e.g, PhenoAge, CausAge, AdaptAge) and immune subsets (**Figure S10f**). In contrast to Factors 8 and 19, its loadings were uniformly aligned with chronological age associations, suggesting that it reflects a molecular axis of gradual, age-like change detectable even within six months (**Figure S10f**, **g**).

Factors 12, 20, and 7, which declined over time (**Figure 5f**), were dominated by saliva metabolites (12, 20) and faecal microbiota (7) (**Figure S10g, h, j**, **Table S7**). Factor 7 correlated with body mass index (**Table S6**), suggesting it primarily captured effects of the intermittent fasting intervention. Factor 12 metabolites were enriched for ammonia recycling pathways (**Figure S10i**), consistent with a shift in nitrogen metabolism. Factor 20 metabolites were enriched for valine, leucine, and isoleucine degradation, phenylalanine and tyrosine metabolism, and the urea cycle (**Figure S10k**). Factors 12 and 20 showed no association with oral microbiota, suggesting their changes were primarily driven by dietary intake or host metabolism changes.

Together, these findings show that measurable biological shifts in immune composition, methylation biomarkers, the faecal microbiome, and oral metabolome occur within just six months, even without a uniform intervention. Some of these shifts tracked with established ageing-associated patterns (e.g., Factor 14, **Figure S10f**), while others reflect subgroup- or intervention-driven dynamics. The results highlight the coexistence of ageing-related and non-ageing influences and underscore the importance of population context when interpreting short-term longitudinal multi-omic changes.

### Ageing-linked features are selectively modifiable by intermittent fasting

The longitudinal multi-omics integration suggested that changes over six months can both resemble or diverge from ageing patterns, but it did not systematically establish which ageing-associated features are themselves modifiable by an intervention. To address this, we examined features associated with chronological age at baseline (FDR<0.05, excluding individual CpGs and ASVs for simplicity). Expected changes were estimated from baseline age-associations (slope × 0.5 years) and compared to the observed longitudinal change between baseline and month 6, stratified by intervention arm and participant compliance. Features were then classified as attenuated, accelerated, or age-opposing (**Figure 6a**).

**Figure 6.**
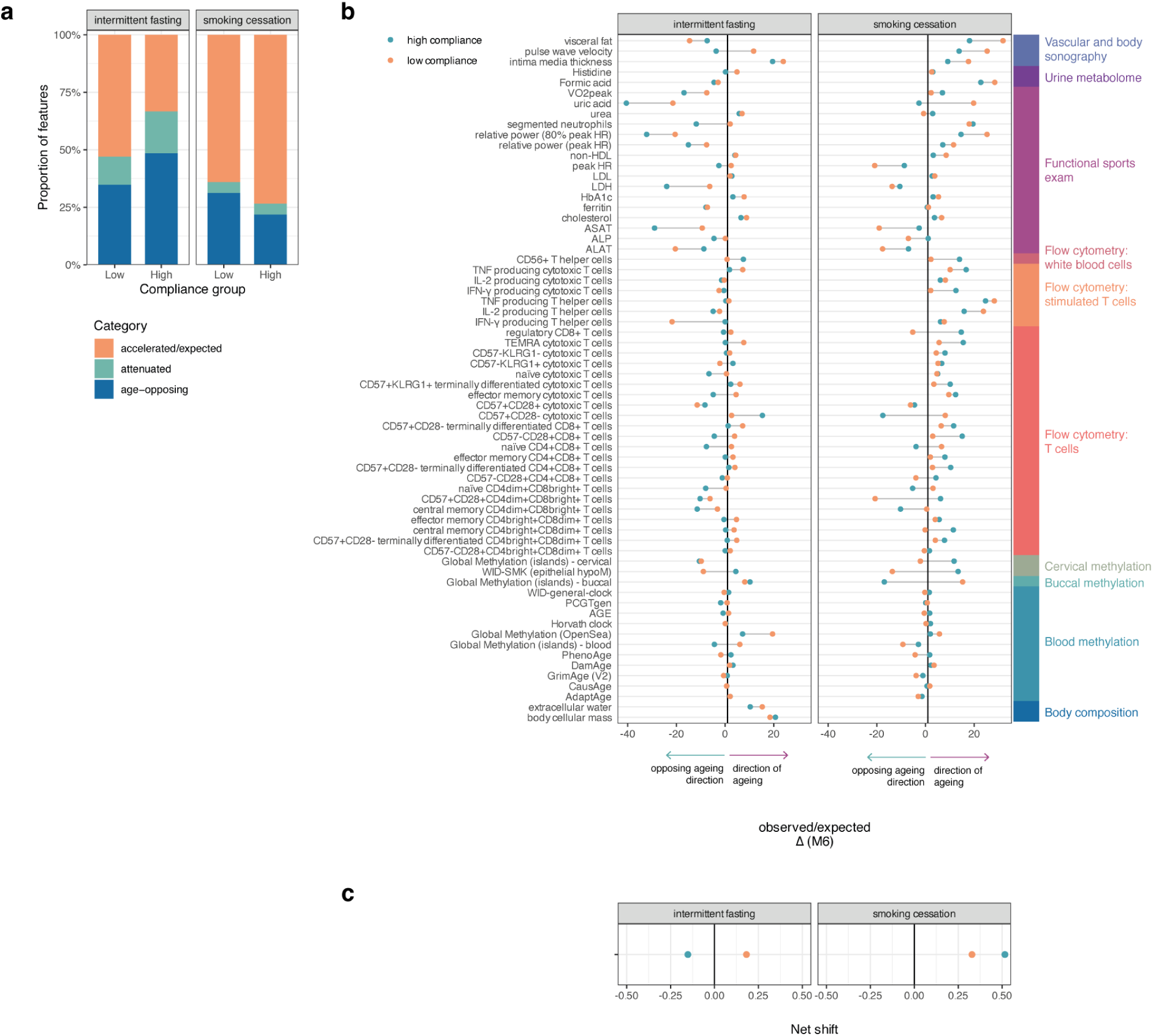
Distinct malleability of age-associated features by intermittent fasting and smoking cessation interventions. **a** Proportion of accelerated-expected, attenuated, and age-opposing changes from baseline to six months by study intervention and compliance group. In the intermittent fasting study arm, compliance was binarised by above or beyond median compliance. In the smoking cessation arm, compliance was binarised based on complete cessation or not. **b** Individual feature ratio changes by study arm and compliance. Features with a significant association with chronological age at baseline were included. **c** Overall net shift by compliance group and study arm.

This analysis revealed striking differences across groups. High compliance with intermittent fasting led to a greater proportion of features shifting in an age-opposing direction, whereas smoking cessation did not show a systematic effect in the same timeframe (**Figure 6a-c**). Among features that changed in the same direction as ageing, highly compliant intermittent fasting participants exhibited attenuated trajectories compared with low-compliance individuals, including vascular intima-media thickness, HbA1c, cholesterol, and extracellular water (**Figure 6b**, **Table S8**). When considering all features jointly, highly compliant intermittent fasting participants displayed a net shift opposite to ageing-associated patterns (**Figure 6c**), consistent with partial reversal of molecular and physiological ageing signatures over six months.

### Blood and mucosal epigenetic biomarkers reveal tissue-specific links to immunity, metabolism, and microbiota

Epigenetic biomarkers or proxies represent powerful tools to quantify ageing ^1^ or disease risk, for instance for women’s cancers ^12,14,16^. However, recent consensus efforts have highlighted limitations in the interpretability and actionability of many ageing biomarkers, hindering their broad clinical translation ^36^. To address this, we leveraged our deeply phenotyped, longitudinal multi-omic dataset to systematically map the molecular correlates of 54 epigenetic biomarkers and proxies using repeated measures correlation (r_rm_) across individuals (**Figure 7**, **Table S9**). As expected, estimated neutrophil proportions from DNA methylation were strongly associated with measured neutrophil counts (r_rm_ = 0.88, p=2.117427e-135, **Figure 7b**, **Figure S11a**), confirming the validity of this proxy. True neutrophil count was also the top correlate for several epigenetic ageing biomarkers, including AgeAccelGrim (V2) (r_rm_=0.52, p=1.9e-29, **Figure 7c**), PhenoAge (r_rm_=0.36, p=9.10e-14), PCGTgen (r_rm_=-0.49, p=2.07e-23), DunedinPACE (r_rm_=0.35, p=5.11e-13), DamAge (r_rm_=0.20, p=4.61e-05) (**Figure S11a**, **Table S9**), verifying that many blood-based methylation biomarkers are sensitive to cell type fluctuations. Blood biomarkers, in particular smoking-related proxies, also reflected variation in other haematologic traits, including haemoglobin (**Figure 6d**) or CRP, suggesting they capture broader immuno-inflammatory states (**Table S9**).

**Figure 7.**
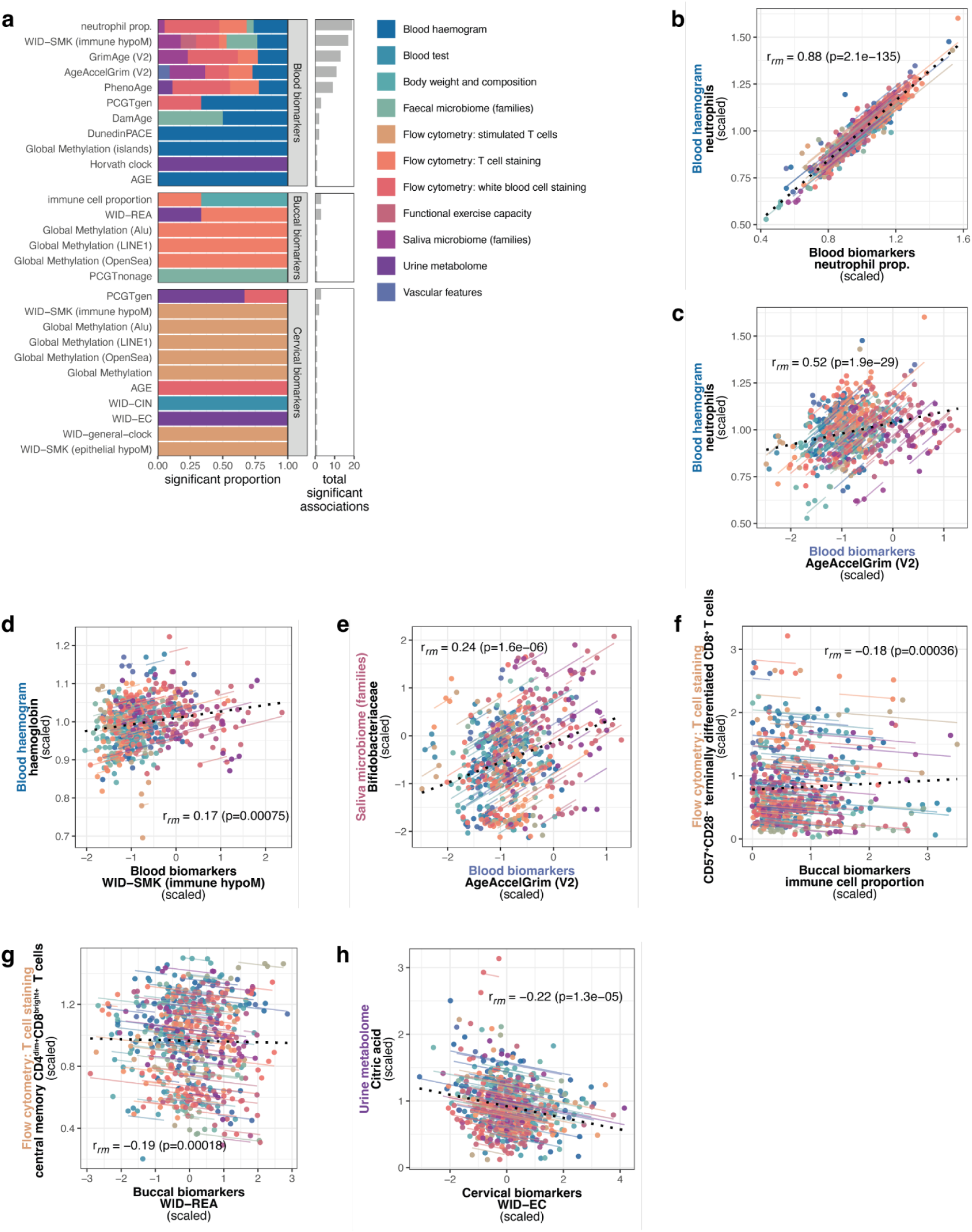
Repeated measures correlation analysis reveals cross-omic association of epigenetic biomarkers and proxies. **a** Overview of the proportion and total significant repeated measures correlations (r_rm_) at fdr < 0.05 per epigenetic biomarker. **b** Repeated measures correlation of the epigenetically-estimated neutrophil proportion and true neutrophil counts in the blood haemogram. **c** Repeated measures correlation of AgeAccelGrim (V2) and true neutrophil counts in the blood haemogram. **d** Repeated measures correlation of WID-SMK immune hypomethylation and haemoglobin values. The directionality of WID-SMK immune hypomethylation has been calibrated so that higher levels indicate higher levels of smoking exposure. **e** Repeated measures correlation of AgeAccelGrim (V2) and salivary *Bifidobacteriaceae* family (CLR-transformed). **f** Repeated measures correlation of buccal immune cell proportion and CD57^+^CD28^-^ terminally differentiated CD8^+^ T cells. **g** Repeated measures correlation of buccal relative-epithelial age (adjusted for immune proportion changes) central memory CD4^dim+^CD8^bright+^ T cells. **h** Repeated measures correlation of the WID-EC index with urinary citric acid. Correlations are obtained from repeated measures correlations (r_rm_). **Abbreviations**: prop., proportion. r_rm_, repeated measures correlation coefficient. CLR, center log ratio.

Several biomarkers showed significant associations with microbiome and metabolite profiles, spanning multiple sample types. DamAge correlated positively with faecal *Clostridia_unk1*, while AgeAccelGrim (V2) was linked to salivary *Bifidobacteriaceae* (**Figure 7e**) and *Mycoplasmataceae*. Although studies have recently begun to explore associations of epigenetic ageing with the intestinal and faecal microbiome^37,38^, the links between oral microbiota and epigenetic ageing are far less explored, representing a novel finding here.

Buccal biomarkers exhibited fewer cross-omic associations than blood biomarkers, and none with blood haemograms. Instead, buccal global repetitive element methylation levels, relative epithelial age, or immune proportion in buccal samples associated with T cell subsets, including central memory CD4^dim+^CD8^bright+^ T cells, CD57^+^CD28^-^ terminally differentiated CD8+ T cells, and naive CD4^dim+^CD8^bright+^ T cells (**Figure 7f, g**, **Figure S11b**), highlighting a surprising link between oral methylation levels and immune subsets. Cervical biomarkers likewise correlated with T cell subsets and selected metabolites (**Figure 7h, Figure S11c**).

These patterns underscore that epigenetic biomarker correlates are strongly shaped by tissue origin, and complementary information may be captured by the application of diverse samples and biomarkers. We note that some associations may only become apparent within specific intervention arms or participant subgroups. These are described in parallel manuscripts focusing on intervention outcomes ^33,35^, providing complementary context to the present cross-omic mapping.

### Epigenome-wide decomposition identifies context-dependent and tissue-restricted CpG plasticity

Understanding whether CpGs are stable, individual-specific, tissue-specific, or intervention-responsive is essential for designing biomarkers that are robust yet sensitive to relevant exposures. In addition to composite biomarkers, the availability of paired, multi-tissue, and longitudinal epigenome-wide data enabled a systematic dissection of the sources of variation at the level of individual CpG sites. We classified CpGs according to their predominant variability component (see STAR Methods). Sites that were stable across all individuals, tissues, and timepoints (n=1,596), and therefore likely of limited functional interest, were excluded from this analysis. Variance partitioning across the remaining sites identified CpG sites that were stable within individuals, tissues, or in interventions, and those exhibiting variability (“malleable”) over time, including those malleable in specific tissues or interventions only (**Figure 8, Table S10**). As expected, inter-individual differences and tissue of origin accounted for the largest proportion of variance (**Figure 8a, b**). Only a minority of CpGs showed predominant variability over time. Among these dynamic sites, most exhibited both tissue- and intervention-specific changes, reflecting localised epigenetic plasticity.

**Figure 8.**
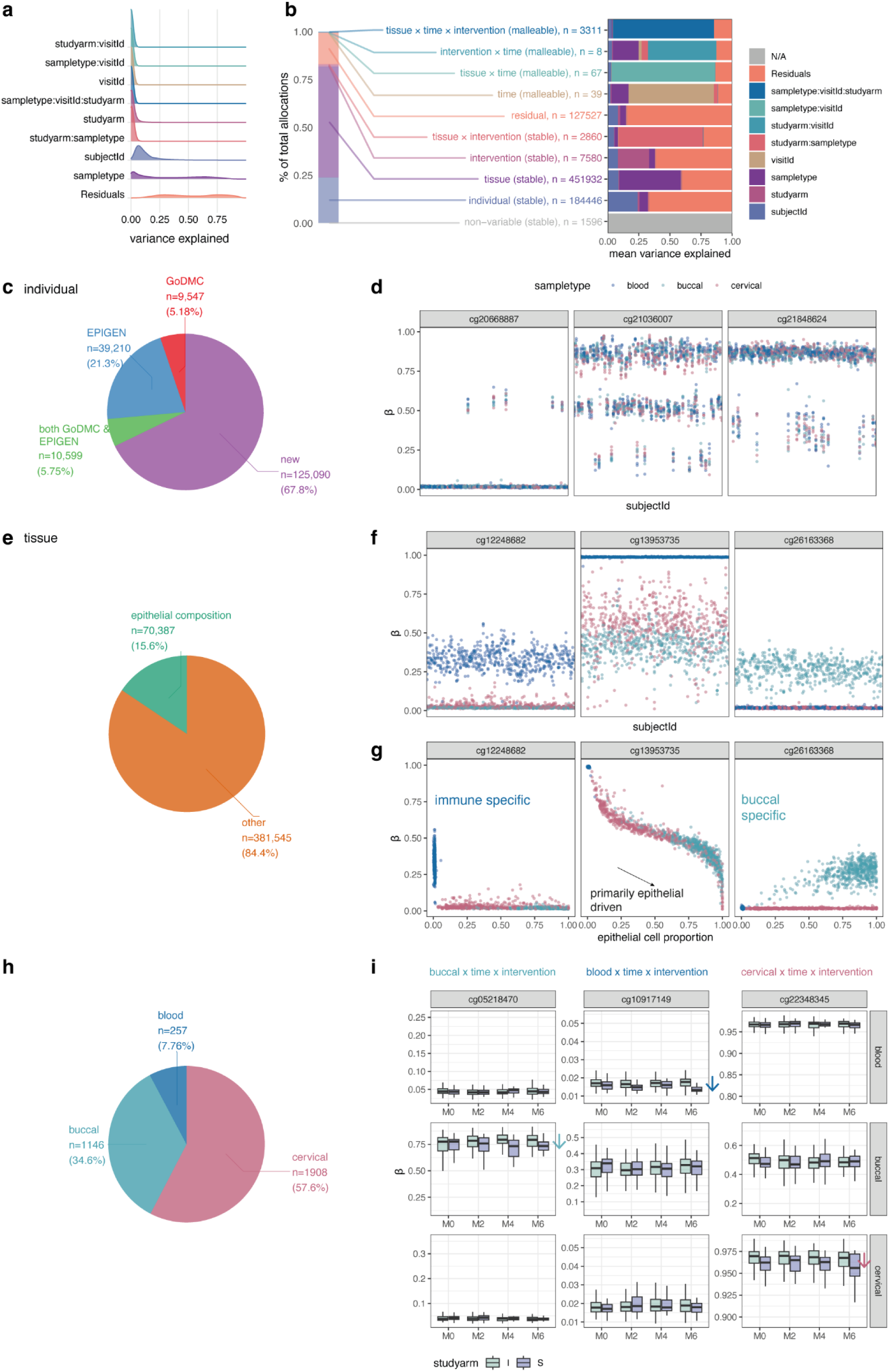
Decomposition of methylation data reveals context-dependent and tissue-restricted CpG plasticity. **a** Proportion of variance explained by model components and residuals. **b** Allocation of CpGs to groups explaining the majority of their variability (left bar chart), and mean variance explained by model components within allocated groups (right panel). **c** Number of CpGs allocated to the individual-variable group that were previously reported as methylation quantitative trait loci (mQTLs) in either GoDMC or EPIGEN, both, or not previously described (new). **d** Visualisation of representative CpG β values in the individual-variable group by subjectId (x axis) and sample type (colour). **e** Number of CpGs in the tissue-variable group that are predominantly explained by changes in epithelial proportion (|correlation|>0.95) or not. **f** Visualisation of representative CpG β values in the tissue-variable group by subjectId (x axis) and sample type (colour). Colour scale for sample types is the same as in panel **d**. **g** Visualisation of representative CpG β values in the tissue-variable group by estimated epithelial cell proportion (x axis) and tissue (colour). Colour scale is the same as in panel **d**. **h** CpGs in the tissue x time x intervention-allocated group variability by tissue. The majority of sites in this group were explained by variability in cervical samples (57.6%), followed by buccal (34.6%) and blood (7.76%). **i** Representative CpGs in the tissue x time x intervention-allocated group by tissue, timepoint, and study arm reveals sites predominantly losing methylation in buccal (*cg05218470*), blood (*cg10917149*), or cervical samples (*cg22348345*) in the smoking cessation group relative to the intermittent fasting group. **Abbreviations**: GoDMC, Genetics of DNA Methylation Consortium. EPIGEN, EPIGEN MeQTL Database. β, methylation beta value. M0-M6, month 0-6. I, intermittent fasting study arm. S, smoking cessation study arm.

Individual-specific CpG sites, a subset of which is shown in **Figure 8d**, were stable within individuals across tissues. Most of the sites identified here have not been previously reported as methylation quantitative trait loci (**Figure 8c**), yet many displayed bi- or tri-modal patterns consistent with underlying genetic variations or SNPs at the probe locus. Among tissue-stable CpG loci, only 16% could be attributed to variation in epithelial cell composition (**Figure 8e**). Tissue-specific sites included features specific to immune cells, epithelial cells, or epithelial cells in a single tissue only (**Figures 8f, g**). We also explored sites variable by intervention and tissue over time, which revealed that the majority of this variance was driven by cervical, followed by buccal and blood samples (**Figure 8h**; examples for each tissue visualised in **Figure 8i**).

This fine-grained decomposition provides a reference for identifying candidate meQTLs, context-dependent biomarkers, and CpGs with potential cross-tissue or longitudinal relevance, offering a framework for interpreting tissue-specific and dynamic methylation changes in future studies.

### Guide to dataset reuse

The analyses above demonstrate the capacity of the dataset to recover known associations, reveal novel cross-tissue links, and decompose molecular variability across biological systems. To maximise the utility of this multi-dimensional resource, we provide an interactive online portal (https://eutops.github.io/lifestyle-atlas) for browsing metadata, accessing harmonised variable dictionaries (SNOMED/SDTM) for cross-study integration, and exploring multi-omic wide association studies (MOWAS). The portal links to raw and processed data deposited in the European Genome-phenome Archive (EGA) and Zenodo, annotated with stable DOIs, and provides documentation for each dataset layer. This resource supports a broad range of applications, including benchmarking of analytical methods, discovery of novel biomarkers, modelling of ageing trajectories, and investigation of system-specific responses to lifestyle interventions. Openly shareable datasets are directly accessible via the portal; restricted-access datasets are available under a data transfer agreement (see Data and Code Availability).

## Discussion

This work presents one of the most comprehensive longitudinal, multi-omic intervention datasets to date, uniquely integrating multi-tissue DNA methylation, metabolomics, microbiome profiles, skin histology and barrier measurements, and detailed immune phenotyping (including functional stimulation assays) in the same individuals. All data were collected under harmonised protocols in a well-characterised cohort, enabling direct, within-cohort comparisons across molecular systems, tissues, and time. This breadth, depth, and integration allow questions to be addressed that are difficult to tackle in more limited datasets, such as the degree to which molecular changes are systemic versus tissue-specific, the relative sensitivity of different modalities to a given exposure, and the consistency of intervention effects with those expected from ageing trajectories.

A particularly notable observation was that features associated with chronological age at baseline did not all follow expected trajectories over six months (**Figure 6a**). In particular. high compliance with intermittent fasting systematically shifted multiple physiological and molecular features in an age-opposing direction, including VO_2_peak, uric acid, ferritin, stimulated immune cell functions, as well as attenuating trajectories of several methylation biomarkers (CausAge, Horvath clock, and others; **Figure 6b**, **Table S8**). This demonstrates that some components of ageing signatures are malleable even over short timeframes, however others appear resistant to modification over a short timeframe. Notably, fewer systematic changes occurred with smoking cessation, in line with different biological effects of these two interventions. Highlighting which ageing-linked features are modifiable by lifestyle change is a key step toward defining actionable biomarkers and surrogate endpoints for healthspan interventions.

Large, well-characterised intervention cohorts have provided invaluable insights into human physiology and ageing. The CALERIE trial, the largest and most detailed caloric restriction intervention in non-obese adults to date (~ 200 participants; 2,327 biospecimens), profiled blood, skeletal muscle, and adipose tissue with whole-genome SNP genotyping and three-timepoint DNA methylation, mRNA, and small RNA data ^21^. While it demonstrated the feasibility of coordinated, longitudinal multi-tissue sampling, it did not include microbiome characterisation, detailed immune phenotyping with functional stimulation, or functional measures such as skin barrier assessments, limiting the ability to analyse cross-system coordination of changes. Other studies have adopted high-dimensional profiling in smaller, targeted cohorts. The NASA Twins Study (n=2) ^39^ integrated multi-tissue genomics, transcriptomics, epigenomics, metabolomics, proteomics, and physiological data to characterise the impact of spaceflight, while a recent exercise intervention study (n=13) ^24^ profiled multiple layers to compare acute and longer-term responses to physical activity. These projects underscore the feasibility and scientific value of multi-omic integration, but are inherently limited in generalisability and power by small sample sizes. Conversely, large-scale observational resources within the Trans-Omics for Precision Medicine (TOPMed) programme, EPIC, TwinsUK, and others, offer complementary breadth and statistical power, with subsets profiled for methylation, microbiome, metabolomics, and other layers. However, these datasets are typically cross-sectional within each omic layer or tissue type and lack longitudinal interventions, making it difficult to dissect malleability or synchronised responses across systems. TirolGESUND bridges these gaps left by combining the molecular depth and multi-tissue scope of intensive small-cohort studies with a larger scale and harmonisation than is typical for such deep interventions. The same participants were profiled across seven tissue and fluid types, multiple omic layers, and repeated timepoints during defined real-world interventions, enabling direct, within-individual comparisons of systemic versus compartment-specific responses. While smaller in size than population-scale cohorts, TirolGESUND captures a broader set of molecular layers, functional readouts, and tissue types than most existing intervention datasets, creating a uniquely integrated platform for exploring cross-system biology in humans.

By analysing this dataset, we highlight several important features of human molecular variability and its responsiveness to lifestyle change. First, variance decomposition across modalities revealed marked contrasts in stability versus responsiveness: clinical phenotypes and immune cell composition were largely stable between individuals, while metabolomics and microbiome features showed high intra-individual variability, consistent with responsiveness to short-term exposures. DNA methylation features, both composite biomarkers and individual CpGs, occupied an intermediate regime, capturing a mixture of stable and responsive states. Second, our multi-omic-wide association study recovered known trait-feature relationships (e.g., age-methylation, smoking-oral microbiota) and identified novel associations, including links between the oral microbiome and menopausal status, or between alcohol intake and skin barrier function. Third, integrative factor analysis captured coordinated cross-tissue molecular signatures for major drivers such as age, smoking, and menopause, as well as less-characterised axes linking oral metabolites, inflammation, and fitness, or alcohol use, extracellular matrix features, and immune variation. Finally, CpG-level decomposition identified context-dependent and tissue-restricted epigenetic plasticity, mapping candidate mQTLs and highlighting the cervix and buccal samples as more dynamic sites for intervention-associated methylation changes than blood.

Beyond these specific findings, the study demonstrates the feasibility and value of combining real-world lifestyle interventions with deep, multi-layer profiling across tissues and time. This design allows us to address questions that are difficult to approach in more limited datasets, such as the degree to which intervention effects are tissue-specific versus systemic, the relative sensitivity of different modalities to a given exposure, and the consistency of molecular trajectories with those expected from chronological ageing. The resource utility is further enhanced by its availability via an interactive portal, providing harmonised variable dictionaries, stable DOIs for datasets, and visualisation tools for exploratory analysis, enabling the community to build on these data for biomarker discovery, benchmarking, and mechanistic studies.

We recognise several limitations of the present dataset. The cohort comprises only women aged 30-60 years from a relatively homogeneous Central European population, which reduces confounding from sex and broad environmental variation but may limit direct generalisability to other demographic groups. The interventions studied here, intermittent fasting and smoking cessation, target distinct biological pathways but do not encompass the full range of lifestyle or pharmacological interventions relevant to healthspan. Sample size, while large for a deeply phenotyped multi-omic intervention study, still limits statistical power for rare features or small-effect associations, particularly when stratifying by intervention arm, adherence, or menopausal status. Additionally, although the longitudinal design captures temporal patterns, follow-up times are relatively short (6 months), although a subset has participated in additional follow-up visits (12, 18 and 24 months). Causal inference remains constrained by the non-randomised, real-world design. Finally, some changes observed over time, particularly those consistent across arms, may reflect general participation effects, unmeasured behaviours, general chronological ageing, or regression to the mean.

Despite these limitations, this work provides a foundational reference for multi-omic, multi-tissue, longitudinal intervention studies in humans. It offers immediate opportunities for the community to benchmark analytical pipelines, validate candidate biomarkers, model cross-tissue coordination, and explore systemic and tissue-specific responses to lifestyle perturbations. The combination of breadth, depth, and harmonisation makes this atlas a valuable platform for advancing biomarker interpretability and actionability, key steps toward the development of reliable surrogate endpoints for healthspan research. Future studies could expand this approach to more diverse populations, intervention types, and longer follow-up, building a network of comparable datasets that together map the multidimensional biology of human ageing and its modulation.

## Resource availability

### Lead contacts

Further information and requests for resources and reagents should be directed to the lead contacts.

### Materials availability

Biological samples are biobanked at the European Translational Oncology Prevention and Screening Institute at the University of Innsbruck. Samples are available upon request by bona fide researchers and access will be decided by a data and sample access committee.

### Data and code availability

All findings supporting this study are available within the main text, Supplementary Methods, and through our data portal (https://eutops.github.io/lifestyle-atlas/). Detailed dataset identifiers, including DOIs and EGA accession numbers, are listed in the STAR methods and accessible via the data portal.

Open-access data are directly linked in the portal and STAR methods. Sensitive datasets, including DNA methylation, health-related data, and microbiome profiles, are deposited under protected access in the European Genome-phenome Archive (EGA) and available to bona fide researchers upon submission of (i) project description and aims, (ii) list of required variables, and (iii) analysis plan. Due to GDPR regulations, these datasets cannot be shared with entities in non-compliant countries. For privacy reasons, direct links between individual-level and multi-omic datasets are not openly provided as the number of variables may increase the risk of reidentification. These can be released only under a data transfer agreement underlying the restrictions for sensitive data. Immune profiling datasets are currently under embargo due to ongoing analyses and will be released upon completion of related studies. Data are available under a CC-BY-NC licence.

Code for data processing and analysis is available at github (<u>github.com/chiaraherzog/MultiOmics_AtlasResource</u>). Code is available under MIT licences.

## Acknowledgements

We thank all volunteers for their participation and commitment to our study.

We thank the clinical and administrative team for their dedication and efforts: Gabriela Hilber, Barbara Rovara, Birgit Kröss, Irene Künzel. We thank Maria Meister, MSc and her team at the Landeskrankenhaus Hall, as well as Christine Kastner and Isma Ishaq-Parveen for their support with processing of samples.

We thank the nutrition and dietetics team, Ute Pichler, Markus Augschöll, David Ebner and Heidi Spildenner who performed dietetic counselling and bioelectric impedance analysis. We thank Dr. Schär AG/SPA for providing the Kanso MCTfiber. We thank Suchthilfe Tirol for their assistance with the dedicated smoking cessation intervention.

We thank the the vascular measurement team Silvia Komarek, Benjamin Dejakum, Alex Messner (all Department of Neurology, Medical University of Innsbruck; VASCage - Research Centre on Vascular Ageing and Stroke, Innsbruck), Bernhard Winder(Department of Vascular Surgery, Feldkirch Hospital, Feldkirch, Austria), and Johannes Nairz (VASCage-Research Centre on Vascular Ageing and Stroke, Innsbruck; Department of Pediatrics II and III, Medical University of Innsbruck).

We thank the student volunteers for their dedication in motivation coaching of participants: Minou Mohraz, Olaya Roces Sanchez, Rosa Huber, Carina Zeitler, Anne Harbring, Rosa Ottmann, Caroline Siebert, Valeria Kovalchuk, Elvira Galeazzo, Lina Förster, Pauline Raßbach, Sonja Breu, Anna Zellmer, and Elif Algül, Melisa Amin, Johanna Haessler, Lilli Schulz, Valerie Nickel, und Lea Rosenstock.

16S rRNA gene amplicon libraries were prepared and sequenced at the Joint Microbiome Facility (JMF) of the Medical University of Vienna and the University of Vienna. We would like to thank Jasmin Schwarz and Joana Séneca Silva for assistance with sample and data processing, respectively.

We thank Prof. Daniela Schmid, Assoc.-Prof. Gaby Sroczynski, and Prof. Uwe Siebert for critical appraisal and feedback of the study protocol.

We thank Dr. Sebastian Schönherr and Lukas Forer for their assistance with the Askimed electronic case report form.

Icons for Figures 1, S5, and the graphical abstract were obtained from Biorender.com.

Chiara Herzog’s current affiliation is King’s College London.

## Author contributions

**Chiara MS Herzog:** Conceptualisation, software, formal analysis, data curation, writing - original draft, supervision, visualisation, project administration.

**Charlotte Vavourakis:** Software, formal analysis, data curation, writing - original draft, visualisation.

**Elisa Redl:** Investigation, supervision, data curation, project administration, writing - review & editing.

**Magdalena Hagen:** Investigation, supervision, writing - review & editing.

**Gabriel Knoll:** Investigation, writing - review & editing.

**Christina Watschinger:** Investigation, supervision, resources, data curation, writing - review & editing.

**Umesh Kumar:** Investigation, writing - review & editing.

**Bente Theeuwes:** Software, writing review & editing.

**Juliane Gasser:** Investigation, writing - review & editing.

**Sophia Zollner-Kiechl:** Investigation, resources, data curation, project administration, writing - review & editing.

**Robert Reihs:** Software.

**Heimo Müller:** Writing - review & editing.

**Maria Cavinato:** Investigation, writing - review & editing.

**Sonja Sturm**: Resources, investigation, supervision, writing - review & editing.

**Hermann Stuppner:** Resources, writing - review & editing.

**Alexander Moschen:** Resources, supervision, writing - review & editing.

**Birgit Weinberger:** Resources, supervision, writing - review & editing.

**Tobias Greitemeyr:** Resources, supervision, writing - review & editing.

**Matthias Schmuth:** Investigation, data curation, project administration, writing - review & editing.

**Verena Moosbrugger-Martinz:** Investigation, writing - review & editing.

**Thomas Trafoier:** Investigation, writing - review & editing.

**Verena Lindner:** Investigation, data curation, writing - review & editing.

**Anna Wimmer:** Investigation, data curation, writing - review & editing.

**Peter Widschwendter:** Investigation, data curation, supervision, writing - review & editing.

**Alexander Höller:** Conceptualisation, investigation, data curation, supervision, writing - review & editing.

**Michael Knoflach:** Investigation, data curation, supervision, writing - review & editing.

**Wolfgang Schobersberger:** Conceptualisation, investigation, data curation, supervision, writing - review & editing.

**Martin Widschwendter:** Conceptualisation, writing - original draft, supervision, project administration, funding acquisition.

## Supplemental information titles and legends

**Supporting information 1. Supplementary figures.**

**Table S1. Included variables and molecular features (excluding CpG-level DNA methylation data).**

**Table S2. Variance decomposition results across omic layers.**

**Table S3. MOWAS analysis results.**

**Table S4. MOFA analysis associations at baseline.**

**Table S5. Top contributors to MOFA factors at baseline.**

**Table S6. Top contributors to longitudinal MEFISTO factors.**

**Table S7. MEFISTO analysis associations.**

**Table S8. Changes over six months with ageing.** Beta age refers to the slope with age, ratio_hi and ratio_lo to the ratio of change in high and low-compliance groups, respectively, with delta_ratio reflecting the difference in ratios. Ratios were categorised into accelerated, attenuated, or age-opposing.

**Table S9. Repeated measures cross-layer correlation analysis for epigenetic biomarkers.**

**Table S10. Methylation variance decomposition results.** Categorisation and variance explained at the CpG-level. See STAR Methods for details. cg, CpG probe name. category, allocated category. dominant, flag if the variance explained for by the category is ≥0.5. Columns starting with varianceExplained_ indicate the proportion of variance explained. residual_flag, flag if the CpG exhibited high levels of residuals. noise_threshold, per-CpG level of technical variance, extracted from replication data. meQTL, indication if CpG is a known meQTL according to GoDMC and/or EPIC or not previously described. celltype_allocation, indicator variable if cell type is primarily driven by epithelial differences (>|0.95|). celltype_type, top correlated cell type with methylation values. celltype_effect, Pearson’s correlation coefficient with top cell type proportion. celltype_pval, p value of correlation with the top cell type. driver_tissue_malleable, indicator of driver of variance for CpGs in the category time × tissue × intervention.

Tables

## STAR ★ Methods

### KEY RESOURCES

### SUBJECT DETAILS

#### Study participants

#### Ethics and study oversight

The TirolGESUND study received ethical approval by the Ethics Committee of the Medical University of Innsbruck (Ethikkommission der Medizinischen Universität Innsbruck #1391/2020, 18.01.2021).

### METHOD DETAILS

#### Recruitment and informed consent

Recruitment occurred from April 2021 until February 2022. Participants were enlisted through targeted recruitment strategies, involving the dissemination of newsletters and flyers among large employers in the Tirol region of Austria, as well as through word of mouth. After an initial screening of registrations of interest, individuals who appeared to fulfill the inclusion criteria were invited to attend webinars, during which comprehensive study information was presented. Following the webinars, participants expressing their interest were invited to the study clinic, where they were provided with sample collection kits. These kits were designed to facilitate the collection of specific samples (urine, saliva, faecal) to be acquired at home, which participants would subsequently bring to the baseline visit. During the baseline visit, participants were once again given an extensive overview of study details and were required to provide informed consent before the study team accepted their home-collected samples. Further samples were obtained at the study clinic, as outlined below.

#### Study setting and visit structure

The study was coordinated from and primarily conducted at the European Translational Oncology Prevention and Screening (EUTOPS) Institute research clinic, located at the Landeskrankenhaus Hall, a regional hospital in Hall in Tirol, Austria. Vascular sonography and optional skin biopsies were conducted at the Departments of Neurology and Dermatology of University Clinic Hospital Innsbruck, respectively. Functional clinical measurements were taken at the Institute of Sports, Alpine Medicine and Health Tourism.

#### Inclusion and exclusion criteria

Participants were allocated to the intervention arm based on inclusion criteria described in the trial registration record (https://clinicaltrials.gov/ct2/show/NCT05678426). Briefly, the inclusion criteria were as follows:

1. Women aged 30 to 60
2. Motivated to change their lifestyle, as indicated by the registration to attend the online webinars for this study
3. Intermittent fasting: non-smoker, BMI between 25 and 35
4. Smoking cessation: ≥10 cigarettes per day for at least the last five years

* If both 3 and 4 applied, the participant was allocated to the smoking cessation group.

Exclusion criteria were:

1. Relevant underlying conditions

a. Current or previous malignant tumour or cancer
b. Current or previous significant cardiovascular disorder; Women with elevated blood pressure were eligible for inclusion on the condition that their blood pressure was effectively controlled under their existing medication regimen
c. Current or previous metabolic disorder (e.g., diabetes type I or II); In the intermittent fasting intervention arm, women with existing hypothyroidism/Morbus Hashimoto were excluded, as the implementation of intermittent fasting may necessitate adjustments to their medication regimen
d. Current or previous psychiatric disorder (e.g., eating disorder, depression)
2. Current pregnancy or lactation period
3. Total hysterectomy
4. Known current or previous premalignant lesion of the cervix uteri (CIN2/3)
5. Concurrent participation in another interventional trial

Participants experiencing any of the exclusion criteria 1b, c, d, 2, 3, and 5 during the study were prospectively defined for exclusion as per the study protocol.

#### Interventions

##### General measures

To enhance motivation and adherence, each participant was assigned a personal coach who provided support and guidance. Regular contact between the participant and coach, with a minimum recommendation of once a week, was encouraged. The coach also maintained records of participant compliance with the main study intervention. Both study arms were offered the opportunity to receive optional adjuvant exercise guidance tailored to their initial fitness levels, but this was not mandatory. Participants were given the option to engage in three supervised exercise sessions throughout the study, focusing on resistance, endurance, and flexibility training.

##### Intermittent fasting intervention

Participants in the intermittent fasting arm followed a “time restricted eating” regime described by de Cabo and Mattson ^40^, guided by a dietician. A shortened stepwise induction was instructed as follows:

- Week 1: 10 h of food intake, 14 h fasting, 5 days per week (2 days habitual diet)
- Week 2: 8 h of food intake, 16 h fasting, 5 days per week (2 days habitual diet)
- Week 3 and subsequent weeks: 8 h of food intake, 16 h fasting for 7 days per week

Optionally, participants could restrict food intake to 6 h per day (18 h of fasting) from week 4 onwards.

Intermittent fasting group participants were randomised to receive a ketogenic supplement (Kanso MCTfiber, medium-chain triglyceride fiber sachet) or not to explore the beneficial effect of ketosis reinforcement, based on its suggested benefits in fasting ^41^. To ensure balanced characteristics in each randomisation arm, we employed menopause- and BMI-stratified block randomisation (blocks of 4). This randomisation process was implemented using a secure Excel worksheet. Participants randomised to the intermittent fasting with ketogenic supplement group were provided with a powdered supplement (Kanso MCTfiber, medium-chain triglyceride fiber sachet), which could be added to food or diluted in water. Recommended use was the use of 1 sachet per day (10 g MCT) during weeks 1 and 2 and two sachets per day (20 g MCT) thereafter for the rest of the study period.

##### Smoking cessation intervention

The smoking cessation arm consisted of three group therapy sessions, accommodating 6-12 participants per session. These sessions were ideally conducted within the first month of the study, with a specific focus on achieving smoking cessation during the second session. Consequently, for each participant, a clearly defined smoking cessation date relative to baseline visit is available (where applicable). Additionally, participants in this arm were offered the choice of two individual telephone appointments with a dedicated addiction counsellor to further support their journey towards quitting smoking.

#### Clinical data collection

##### Electronic case report form

Data were recorded electronically in an electronic case report form (eCRF), Askimed (https://www.askimed.com/, 1.21.3) which included data plausibility and validation checks, or in the Qualtrics survey environment (baseline epidemiological questionnaire only) (April 2021; https://www.qualtrics.com).

##### Functional clinical measurements

Functional clinical evaluation included ergometry, spirometry, and blood chemistry taken at baseline and month 6. Measurements were collected at the Institute for Sports Medicine, Alpine Medicine and Health Tourism. Values were recorded in the eCRF and automatic data validation was performed using previously defined plausible ranges. All participants performed an incremental cycle ergometry test to exhaustion, starting at 50W with an increase of 25W for every 90s. VO_2_peak was calculated according to the following equation:

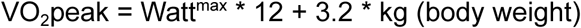

##### Vascular assessment

Vascular measurements included measurement of pulse wave velocity and plaque score (baseline and month 6). Measurements were obtained at the Department of Neurology at the Medical University of Innsbruck. Values were recorded in the eCRF and automatic data validation was performed using previously defined plausible ranges.

##### Body composition measurements

Height and weight of participants were recorded at each visit for calculation of the body mass index (BMI). Bioelectric impedance measurements using the BIACORPUS RX4000® device were taken by a trained dietician or nurse from participants in the intermittent fasting arm at each study visit. Prior guidelines ^42^ were followed to ensure uniform measurements, including standardisation of last meal and exercise times. Briefly, the participant was asked to rest in a horizontal position, with no contact between limbs and torso. The skin was disinfected and cleaned. Electrodes were placed in the correct position and the BIACORPUS RX4000® device was switched on to obtain measurements. Measurements were recorded in the eCRF and automatic data validation was performed using previously defined plausible ranges. Quantification of abdominal and subcutaneous fat was conducted at baseline and month 6 the Department of Neurology at the University Clinic Hospital Innsbruck, adapting a previous protocol ^43^. Triplicate measures were taken. Values were recorded in the eCRF and automatic data validation was performed using previously defined plausible ranges.

##### Compliance data

Compliance with the intervention was documented in the eCRF by personal coaches based on participant reports during weekly contact and study visits conducted by doctors. Coaches recorded the number of compliant days per week in the intermittent fasting arm, defined as the number of days out of 7 the participant adhered to the fasting intervention (typically 16 h fasting per day or more). In the smoking cessation arm, the average daily cigarette consumption was recorded as well as the number of smoke-free days each week. Nicotine replacement was also queried, including nicotine gum, plaster, and the use of electronic cigarettes or heated tobacco products. Study doctors provided unstructured comments on compliance and participant feedback, which were later standardised and harmonised with coach data to derive a compliance score (weighted compliant days per week, defined as the % of days in the entire study period (baseline to month 6) with 16 h fasting or smoke-free days, respectively, over the duration of the study). Supplement use was also queried in an unstructured manner by doctors and coaches. Supplement use was not included in the compliance score but considered for further analysis.

##### Participant questionnaires

Participant questionnaires for the collection of epidemiological data and medical history (custom questionnaire), quality of life (EQ-5D-5L) ^44^, and exercise frequency data (International Physical Activity Questionnaire, IPAQ) ^45^ were conducted online via secure links and a password-protected login provided to participants.

##### Data harmonisation

Data from the eCRF and Qualtrics survey platform were extracted and harmonised by the lead investigator. To ensure interoperability, look-up tables using machine-readable, standardised terminology identifiers from Systematized Nomenclature of Medicine-Clinical Terms (SNOMED-CT) Identifier (SCTID) and Study Data Tabulation Model (SDTM) terminology (where available) were generated and are available on Zenodo (doi to be released upon publication).

#### Sample collection and processing

Sample collection, processing, and downstream analyses followed predefined standard operating procedures.

##### Buccal sample

To ensure adequate DNA yield for molecular profiling of buccal cell samples, we utilised four Qiagen OmniSwab devices (#WB100035), with two devices used per side. The cheeks were swabbed with external counterpressure and swab heads were placed in a single tube and stored at −20°C until DNA extraction for subsequent epigenetic analysis.

##### Blood sample

Blood was drawn by trained personnel and collected into various devices depending on intended use.

##### Cervical sample

Samples were collected during the study visit using a speculum. The brush was rotated five times and placed in the Hologic ThinPrep® collection device. The device was transported to the lab and samples were pelleted. The pellet was washed once using phosphate buffered saline, and subsequently stored at −80°C until DNA extraction for subsequent epigenetic analysis.

##### Saliva sample

Participants were asked to collect two saliva samples at home on the day of the study visit: native and DNA/RNA stabilised. Participants were asked to not eat, drink, smoke, or use oral hygiene products for one hour prior to sample collection, and ideally collect samples prior to brushing teeth. For the native sample, the mouth was rinsed with water and 5 minutes later, 2 mL of saliva were placed in a collection tube. The sample was placed in the provided cooling bag along a freezer ice pack until the study visit, where it was aliquoted and stored at −80°C until further processing. The stabilised sample was placed into a Zymo Research DNA/RNA Shield Saliva Collection Kit tube (#R1210-E) and inverted 10 times. At the study visit, it was aliquoted and stored at −20°C until further processing.

##### Urine sample

Participants were asked to collect a urine sample at home on the day of the study visit. Participants collected morning urine and placed the collected sample in the cooling bag alongside a freezer ice pack until the study visit, where it was centrifuged to separate the supernatant and pellet. The supernatant was aliquoted and stored at −80°C until further processing. The pellet is stored at −80°C.

##### Faecal sample

Participants were asked to collect two faecal samples at home as close as possible to the day of the study visit: native and DNA/RNA stabilised. The native sample was placed in the appropriate tube and placed in the provided cooling bag alongside a freezer ice pack until the study visit, where it was aliquoted and stored at −80°C until further processing. The stabilised sample was placed in a DNA stabilisation solution (Invitek Stool Collection Tubes with DNA Stabilizer, #1038111), stored at −20°C until further processing.

##### Skin biopsy and transepidermal water loss assay

Participants were invited to provide optional skin biopsies at baseline and 6 months. Biopsies required a separate informed consent. Two to three 6 biopsies were taken from the skin of the inner arm after disinfection and local anaesthesia. Tissue samples were transferred to sample vials, and placed in liquid nitrogen or formaldehyde for subsequent analysis. The biopsy site was sutured, with sutures removed after 10-14 days. Participants were provided with detailed instructions on post-biopsy skin care.

To measure epithelial barrier function, an independent and objective parameter of epithelial differentiation ^46,47^, we conducted a non-invasive transepidermal water loss (TEWL) assay. TEWL was measured on the volar forearm using a Tewameter (Courage & Khazaka, Cologne, Germany), under comparable environmental conditions ^48,49^. Participants were advised to not use any topical medications or emollients one week prior to measurements. Exclusion criteria included active skin disease; i.e., patients with skin conditions that are known to be associated with a barrier defect including eczema, atopic dermatitis, psoriasis, ichthyosis. The maximal value within a 30 s assessment period was used for subsequent analyses.

#### Wearable device data

Participants were provided with a wearable fitness tracker (Garmin® vivosmart 4) to capture basic activity levels (daily steps, active kilocalories, minutes of vigorous intensity per day), basal metabolic rate, resting heart rate, sleep behaviour (REM sleep duration), and a proprietary measure termed fitness age. Data was accessed via the Garmin Health API. To evaluate levels of dietary ketosis, a subset of participants in the intermittent fasting study arm received a device to measure capillary ketone bodies (FORA 6) and ketone body strips. Participants were encouraged to regularly measure β-hydroxybutyrate levels (2-3 times per week) at the end of the fasting period. Data was accessed via the FORA Telehealth Option.

#### Molecular assays

##### Blood counts and chemistry

Routine blood counts and chemistry were assayed at each visit from four mL of peripheral venous blood collected in LiHeparin/EDTA tubes by trained staff.

##### DNA methylation data

DNA from blood was isolated according to the blood protocol of the Mag-Bind® Blood & Tissue DNA 96 Kit (#M6399-01, Omega Bio-tek) with minor adjustments (560 µl of HDQ binding buffer, 20 µl of beads). DNA from cervical samples was isolated using the cells and tissue protocol of the Mag-Bind® Blood & Tissue DNA 96 Kit (#M6399-01, Omega Bio-tek) kit with an additional SPM washing buffer step. DNA from buccal swabs was isolated using the QIAamp DNA investigator kit (#56504, Qiagen) according to the manufacturer’s protocol. All DNA samples were quantified using the Quantifluor® dsDNA System (#E2670, Promega). Bisulfite modification of 250 ng was carried out using the EZ DNA Methylation-Lightning kit (#D5030, Zymo) on a Tecan Fluent 480 liquid handling platform and DNA was standardised to a concentration of 21 ng/µL bisulfite-converted DNA. Eight µl of bisulfite-converted DNA were processed on the Illumina Human MethylationEPIC v1.0 array (#20042130) according to the manufacturer’s instructions. To minimise batch effects across longitudinal analyses from the same individuals, samples derived from the same participant were kept on the same beadchip whenever possible, with positions within the beadchip randomised. Empty spaces were filled with technical replicates of a random selection of samples (n=15) to evaluate reliability of methylation signals.

##### Faecal DNA extraction for microbiome profiling

DNA isolation for microbiome sequencing from faecal samples was performed using the Invitek PSP® Spin Stool Basic Kit (#1038120300). DNA isolation was carried out according to the manufacturer’s protocol with minor modifications. Specifically, after the washing step with Wash Buffer II (step 7 of the protocol), an additional washing step with 700 µl EtOH absolute was interpolated. Samples were eluted in 50 µL of Elution Buffer (step 9). DNA concentration was measured using a NanoDrop™ device. When necessary, DNA was further purified with Invitek MSB® Spin PCRapace Kit (#1020220300).

##### Saliva DNA extraction for microbiome profiling

DNA for microbiome sequencing was isolated from saliva samples with the corresponding Zymo Research ZymoBIOMICS™ DNA Miniprep Kit #D4300) according to the manufacturer’s protocol, with several modifications: in step 1, 1000 µL of stabilised saliva were used for DNA extraction. Samples were transferred into ZR BashingBead™ Lysis Tubes, supplemented with 20 µL Proteinase K, and shaken on an Eppendorf® ThermoMixer® C (#5382000015) at 900 rpm for 40 minutes at 55°C. In step 2, samples were homogenised using a Vortex-Genie® 2 Mixer (Scientific Industries, #SI-0236), adapted to a horizontal-(24) microtube holder (Scientific Industries, #SI-H524) for 40 min at maximum speed. In steps 3, 8, 9 and 10, the centrifugation speed was increased to 16,000 rcf for 1 min. An additional washing step was performed after step 10 with 700 µL EtOH absolute with subsequent centrifugation at 16,000 rcf for 1 min. At step 11, DNA was eluted in 55 µL of ZymoBIOMICS™ DNase/RNase Free Water, which was preheated to 60°C. DNA concentration was quantified using a NanoDrop™ and where necessary, DNA was further purified using the Zymo Research Genomic DNA Clean & Concentrator®-10 kit (#ZD4011).

##### Microbiome sequencing 16S library generation

Library generation, 16S rRNA gene amplicon sequencing, inference of Amplicon Sequence Variants (ASVs) and taxonomic classification was done at the Joint microbiome facility (JMF) in Vienna. Briefly, amplicons of the variable V4 region (primer pair 515F Parada ^50^-806 Apprill ^51^) of the 16S rRNA gene were generated using dual barcoding in a two-step PCR approach ^52^ and sequenced on an Illumina MiSeq platform.

##### Nuclear magnetic resonance (NMR) analysis for urine and saliva metabolites

Frozen urine and saliva samples stored at −80°C were thawed at room temperature and solid particles were sedimented at 14,000 rpm for 10 min. 540 µL urine and saliva supernatant were taken and added to 60 µl of a buffer containing KH2PO4 (1.5 M), NaN3 (2 mM) and 0.1% 347 TSP in 100 mL D2O at a pH of 7.4. After brief vortex mixing and centrifugation at 10,000 rpm for 5 min, then 600 µL of the samples were transferred to NMR tubes and analysed directly. NMR spectra were acquired using an Avance II 600 MHz 349 spectrometer (Bruker Biospin, Rheinstetten, Germany), equipped with a Bruker 5 mm CryoProbe Prodigy TCI probe head with Z-gradient. Samples were measured in automation using a Bruker BACS-60 sample changer operated by IconNMR and Topspin 3.2 (Bruker Biospin). The probe head temperature was adjusted to 300.0±0.1 K using a 99.8% deuterated methanol sample, and water suppression was calibrated using a sample containing 2 mM sucrose, 0.5 mM TSP, and 2 mM NaN3 in 90% D_2_O. Prior to measurement temperature of each sample was equilibrated to 300.0±0.1 K for 5 min and automatic tuning and matching, locking to D2O signal, shimming, and pulse calibration were performed. For each sample, a 1D 1H-NOESY with water suppression via pre-saturation, 32 scans, a receiver gain of 90.5, a relaxation delay d1 of 4.0 s, a mixing time d8 of 0.01 s, and an acquisition time of 2.726 s was acquired. Additionally, using standard pulse sequences and experimental settings, 1H–J-Res, COSY, HMBC and HSQC-spectra were acquired for peak annotation and substance identification in selected samples.

##### Peripheral mononuclear cell isolation

Isolation of peripheral mononuclear cells (PBMCs) from blood was performed by density gradient centrifugation using the density gradient medium Lymphoprep (STEMCELL Technologies, #07851), within 24 h of sample collection. Samples were stored at room temperature until PBMC isolation. Isolated PBMCs were frozen in fetal calf serum (FCS; Sigma-Aldrich, #F7524-500) containing 20% dimethyl sulfoxide (DMSO; Sigma-Aldrich, #D2650) and stored in liquid nitrogen.

##### Cell culture for immune cell stimulation

Cryopreserved PBMCs were thawed in prewarmed RPMI 1640 medium (Sigma-Aldrich, #R8758-6×1L) containing 20% FCS (Sigma, #F7524-500), 1% P/S (Sigma-Aldrich, #P0781-100ML) and 2.5% HEPES (Sigma, #H0887). Free DNA was digested with DNase I (Roche, #11284932001) for 1h at 37°C and PBMCs were subsequently incubated overnight in RPMI 1640 with 10% FCS at 37°C. To analyse cytokine production of CD3^+^CD4^+^ and CD3^+^CD8^+^ T cells, rested PBMCs were stimulated for 16 h at 37°C with 1 µg/mL α-CD28 (BioLegend, Cat #302934, Clone CD28.2) and 1 µg/mL α-CD3 (BioLegend, Cat #317326, Clone OKT3) in the presence of 10 µg/mL Brefeldin A (BFA; Sigma, #B6542-5MG).

##### Flow cytometry profiling

Immunofluorescence surface staining was performed by adding a panel of directly conjugated antibodies to PBMCs. Dead cells were excluded from the analysis using a fixable viability stain (BD, #564997). For the detection of cytokine production after surface staining, cells were permeabilised using the Cytofix/Cytoperm Fixation/Permeabilization Kit (BD, #554714) and incubated with intracellular antibodies. To detect intracellular FoxP3 expression, surface stained PBMCs were permeabilised using the Transcription Factor Buffer Set (BD, #562574) and incubated with α-FOXP3. Labelled cells were measured by an LSRFortessa Cell Analyzer (BD). Data were analysed using Flowjo v10 software. A table of antibodies is provided in the STAR Methods Details.

##### Skin immunohistochemistry

Biopsies were washed in AA-PBS (PBS containing 250 µg/mL Amphotericin B, 10.00 IE (µL)/mL Penicillin/Streptomycin) and immersed in 4% Paraformaldehyde fixative solution for 24 h. Biopsies were further processed for embedding in paraffin blocks. 5 µm serial sections were obtained and used for Hematoxylin/Eosin (H&E) staining or for immunohistochemistry for the detection of senescence-related proteins. H&E-stained slides were used for morphometric analysis for the following parameters: epidermal thickness, thickness of the stratum corneum, number of epidermal cells per field. The measurements were standardized using millimetric scale plate (Carl Zeiss®) for the several objectives used (4, 10, 20X). This procedure was performed for at least 20 epidermal measures from each sample.

### QUANTIFICATION AND STATISTICAL ANALYSIS

#### Preprocessing pipelines

##### Methylation data preprocessing

Methylation beta matrices were prepared from raw .idat using a previously published ^13,53^, standardised preprocessing pipeline suitable for diverse sample types, eutopsQC (version 1.2, https://github.com/chiaraherzog/eutopsQC). Briefly, raw data were loaded using the R package minfi ^54^, version 1.55.1, and samples with median methylated and unmethylated intensities <9.5 or with >10% failed probes were removed. Background intensity correction and dye bias correction were performed using the minfi single sample *preprocessNoob* function. Probe bias correction was performed using the beta mixture quantile normalisation (BMIQ) algorithm implemented in the ChAMP package ^55^, version 2.39.0. Probes that mapped to the Y chromosome, previously flagged SNPs ^13,54,56^, non-CpG probes, and any probes with >10% failure rate were removed. Beta values from failed probes were imputed using the impute.knn function as part of the impute R package. The fraction of immune cell contamination, and the relative proportions of different immune cell subtypes in each sample, were estimated using the EpiDISH algorithm ^57^, version 2.25.0, using the epithelial, fibroblast, and immune cell reference dataset (centEpiFibIC.m). For more granular interpretation of cell type proportions, the hierarchical EpiDISH algorithm was applied, with the secondary reference matrix cent12CT.m. The top 30,000 most variable probes (ranked by standard deviation) were used in a principal component analysis. We explored associations of the top 10 principal components with biological and technical features using correlation analysis (for continuous covariates) or Kruskall-Wallis tests (for categorical variables). Samples from matched individuals were evaluated using SNP control probes present on the Illumina array. Mismatches were investigated and, where possible, samples assigned to the correct individual post-hoc. Methylation biomarkers were computed using previously published algorithms or weights according to authors’ guidelines and verified using the biolearn package ^58^ (**Figure S3**). Composite cervical and buccal methylation biomarkers were residualised for estimated immune cell proportion to account for fluctuations in major cell types.

##### Microbiome data preprocessing and QC

For each sequence run, pooled amplicon libraries were demultiplexed with the python package demultiplex (Laros JFJ, github.com/jfjlaros/demultiplex). PhiX contamination was removed, and barcodes, linkers and primers were trimmed using BBduk (BBTools, Bushnell B, sourceforge.net/projects/bbmap). Estimation of the sequence error rates, read abundance calculation, merging of read pairs, and sample ASV inference was done with the R package DADA2 ^59^, as recommended in a community guideline ^60^. Initial QC was done after merging results from separate sequence runs for each sample type, i.e., stool and saliva. Taxonomic classification of the ASVs was done against the SILVA database SSU Ref NR 99 release 138.1 with the SINA classifier, version 1.7.2 ^61^. Singleton and doubleton ASVs, ASVs classified as eukaryotes, mitochondria or chloroplasts, and those resulting from known wet-lab contaminants were removed. Rarefaction curves and species (ASV) richness were calculated with the rarecurve() and rarefy() functions in the R package vegan (version 2.6-4, https://CRAN.R-project.org/package=vegan). Microbiome data (family and species-level) were center log ratio (CLR)-transformed.

##### Metabolome data preprocessing

Raw data underwent automatic Fourier transformation, phase correction, and baseline correction. Topspin 3.2 (Bruker) was employed for data processing, while Chenomx Profiler (Chenomx Inc., version 8.1) and the Human Metabolome Database HMDB (Version 5.0) ^62^ were utilised for the annotation of NMR peaks to compounds. This comprehensive approach ensures accurate and reliable processing of NMR data, allowing for the extraction of meaningful information from complex spectra. The spectral concentration was converted into comma-separated values (CSV) using the Microsoft Excel format for multivariate statistical analysis.

#### Analyses

##### Simple variance partition

We initially explored variance between individuals, within individuals, and by intervention/study arm leveraging linear mixed effects models implemented in the R variancePartition package ^63^, version 1.39.0. The formula applied was ~ (1 | participant) + (1 | study arm), whereby variance explained by participant was classified as inter-individual variation, variance explained by study arm allocation was classified as intervention/study arm variation, and residual variance was classified as intra-individual variation, including variation over time and other unmeasured effects. For probe-level simple DNA methylation variance partition, we included only more reliable DNA methylation probes in each sample type (blood, buccal, and cervical), leveraging an approach developed by Nazarenko et al ^26^. We estimated unreliability and median intensity across technical replicates for each tissue type and retained the top 10% of reliable probes, leaving approximately 78,000 CpG sites.

##### Multi-omic-wide association studies (MOWAS)

Multi-omic-wide associations between chronological age, menopause, BMI, current or ever smoking, VO_2_peak, and alcohol use were computed leveraging baseline blood haemogram, skin histology and transepidermal water loss assay, functional sports exam, vascular and body sonography, flow cytometry, metabolome (urine and saliva), microbiome families and ASVs (faecal and saliva; CLR-transformed), and methylation data, using linear models (omic feature ~ variable). Menopause, BMI, current or ever smoking and VO_2_peak were adjusted for chronological age, while alcohol use was adjusted for both chronological age and smoking. Methylation sites (CpGs) were prefiltered to improve efficiency in subsequent linear models, retaining only those with a Pearson correlation estimate >|0.1|. For methylation features, models were adjusted for features noted above, as well as cell type proportions (granulocyte proportion, i.e., sum of monocytes, neutrophils and eosinophils for blood, immune cell proportion for cervical and buccal samples). Features significant at Bonferroni-corrected p values <0.05 were considered significantly associated with the phenotype, with the exception of menopause and VO2_peak_, where features at FDR <0.1 were considered significant. Multiple testing correction was applied within each data type and phenotype.

##### Multi-omic factor analysis (MOFA) and Integration of temporal omics data using MEFISTO (Method for the Functional Integration of Spatial and Temporal Omics data)

MOFA^29^ and its longitudinal extension, MEFISTO^32^, were conducted using the R package MOFA2, version 1.13.0. Layers with fewer than 15 features were omitted (e.g., Flow cytometry: T cell stimulation). Normalised, centred data were used for MOFA and MEFISTO.

A total of 632 features were used in MOFA across 12 layers: blood methylation biomarkers, n=17; buccal methylation biomarkers, n=15; cervical methylation biomarkers, n=19; flow cytometry (T cell staining), n=26; flow cytometry (white blood cell staining), n=27; saliva microbiome families, n=52; faecal microbiome families, n=76; saliva metabolome, n=49; urine metabolome, n=36; blood methylation PCs, n=132; buccal methylation PCs, n=112; cervical methylation PCs, n=71. Hyperparameters were optimised to explore the optimal number of factors. A total of 10 factors were included in the final baseline model.

For MEFISTO, a total of 1,032 features were used in MOFA across 12 layers: blood methylation biomarkers, n=17; buccal methylation biomarkers, n=15; cervical methylation biomarkers, n=19; flow cytometry (T cell staining), n=65; flow cytometry (white blood cell staining), n=30; saliva microbiome families, n=52; faecal microbiome families, n=76; saliva metabolome, n=49; urine metabolome, n=36; blood methylation PCs, n=327; buccal methylation PCs, n=237; cervical methylation PCs, n=109. Time was modelled continuously (from days since baseline). For the overall MEFISTO model, 20 factors gave the best result.

We assessed associations of factor values with phenotypic features through both linear/logistic regression and Spearman correlation/Kruskal-Wallis tests (for continuous or numeric and categorical variables, respectively). P values were FDR-corrected for multiple testing. Phenotypic features with q values <0.05 for both the regression and alternative testing strategy were considered significant.

##### Assessment of ageing-associated feature malleability

To evaluate whether ageing-associated features were modifiable by intervention, we retained features associated with chronological age at baseline at FDR<0.05, excluding individual CpG sites and ASVs. For each age-associated feature, we then estimated the expected six-month change under ageing by multiplying the baseline age-effect slope (beta_age) by the study interval (0.5 years). Observed longitudinal changes (month 6 vs. baseline) were aligned to the ageing direction and compared against the expected magnitude. Each feature was classified into one of four categories: age-opposing (observed change opposite in sign to the expected ageing effect), attenuated (change in the same direction but smaller than expected), unchanged (change consistent with the expected magnitude; ±10% tolerance), and accelerated (change in the same direction but larger than expected). Expected and observed changes were computed separately for each intervention arm (intermittent fasting, smoking cessation) and stratified by compliance (high vs. low, defined by median split of compliance score for fasting, or complete cessation vs. continued smoking). To compare a net shift, we summarised results across features to quantify the net direction of ageing modification in each intervention and compliance group, assigning −1 to age-opposing, +1 to accelerated, and 0 to unchanged or attenuated features, and averaging across all features at M6.

##### Repeated measures correlation analysis

Repeated measures correlation analysis was conducted using the R rmcorr package, version 0.7.0. In brief, 54 epigenetic biomarkers or proxies across buccal, cervical, and blood samples were selected for this analysis. Epigenetic biomarkers in buccal and cervical samples were adjusted for immune cell proportion using baseline methylation samples (i.e., leveraging residuals after correcting for the association of baseline values with immune cell proportion, as estimated by EpiDISH). Repeated measures correlation of each biomarker with cross-omic variables were run using scaled data. Findings were FDR-corrected for each biomarker, retaining associations at FDR<0.05.

##### Detailed methylation variance partition

To annotate CpGs with their primary source of variance, we implemented an expanded variance decomposition using the R variancePartition package, version 1.39.0. We initially decomposed the variance of methylation M values according to the following formula: ~ (1|subjectId) + (1|sampletype) + (1|visitId) + (1|studyarm) + (1|studyarm:visitId) + (1|studyarm:sampletype) + (1 | sampletype:visitId) + (1|sampletype:visitId:studyarm), allowing us to decompose variance proportions into those explained by individual (subjectId), sample type (sampletype), time (visitId), intervention (studyarm), intervention over time (studyarm:visitId), intervention by tissue (studarm:sampletype), tissue by time (sampletype:visitId), and the interaction between tissue, time, and intervention (sampletype:visitId:studyarm), returning the proportion of variance explained by each of the above, as well as residual variance (not explained by the above). The model was run over each CpG, subsetting those present in both the technical replicate and full dataset after quality control (n=779,366). CpGs whose overall variability was below the variability of technical replicates (n=1,596) were excluded from variance decomposition.

We then classified each CpG according to the main source of variance explained in the variance partitioning analysis, considering eight possible components: tissue × time × intervention; tissue × time; intervention × time; time; tissue × intervention; intervention; individual; and tissue. We assigned each feature to the component explaining the largest fraction of variance, provided this fraction exceeded a CpG-specific threshold derived from technical variance (i.e., noise floor estimated from technical repeats and mapped to a per-CpG variance explained cutoff). CpGs with no component above this technical threshold were labelled as residual. We further annotated whether the top component was dominant (>0.5 of variance) or non-dominant (<0.5). Components involving a time term were labelled malleable (tissue × time × intervention; tissue × time; intervention × time; time), all others were stable. We also flagged CpGs with high unexplained variance (residuals > 0.5).

##### Methylation quantitative trait loci analysis

To explore individual-specific CpGs further, we obtained information on known methylation quantitative trait loci (meQTLs) from GoDMC^64^ (450K array; n=85,568 unique CpGs) and EPIGEN meQT^65^ (EPIC array, n=247,429 unique CpGs) databases and overlapped them with CpG sites in our dataset.

##### Tissue-specific CpGs

To explore tissue-specific CpGs further, we explored each CpG’s correlation with one of 14 cell types (epithelial, fibroblast, CD4 naive T cells, basophils, CD4 memory T cells, memory B cells, naive B cells, regulatory T cells, memory CD8 T cells, naive CD8 T cells, eosinophils, NK cells, neutrophils, and monocytes). Full data are provided in **Table S9**. CpGs whose correlation with epithelial cell proportion was greater than |0.95| were considered to be primarily driven by changes in epithelial cell proportions.

##### Exploration of CpGs with intervention-specific malleability

To identify the tissue most strongly contributing to tissue × time × intervention variability we performed a screening analysis on CpGs previously classified as dominated by this triple interaction. For each tissue, we computed the variance across intervention × time group means, using M-values restricted to samples from that tissue. Group means were calculated for each combination of intervention arm and timepoint, and the variance across means was computed per CpG. The tissue with the highest group-mean variance was assigned as the “driver tissue” for that CpG.

## DATA AND CODE AVAILABILITY

Code to reproduce analyses is provided on Github ((https://github.com/chiaraherzog/MultiOmics_AtlasResource).

