## Supporting information for "Multi-modal atlas of lifestyle interventions reveals malleability of ageing-linked molecular features"

**Figure S1. Overview of the TirolGESUND participant flow.** **2**
**Figure S2. Technical validation of methylation data.** **3**
**Figure S3. Technical validation of external biomarker value algorithms.** **5**
**Figure S4. Technical validation of metabolomics and microbiome data.** **6**
**Figure S5. Data structure overview.** **7**
**Figure S6. Association of latent factors 1 and 2 deduced from a MOFA analysis at baseline with smoking, body-mass index, age, and intima-media thickness.** **8**
**Figure S7. Baseline MOFA reveals menopause-associated immune remodelling.** **9**
**Figure S8. Oral microbiota at baseline are associated with immune infiltration while oral metabolites associate with systemic inflammation and exercise.** **10**
**Figure S9. Microbiome and immune features associated with smoking, menopause, and skin barrier.** **11**
**Figure S10. Global changes in immune cells, ageing features, faecal microbiome, and oral metabolome.** **12**
**Figure S11. Repeated measures correlation overview.** **14**

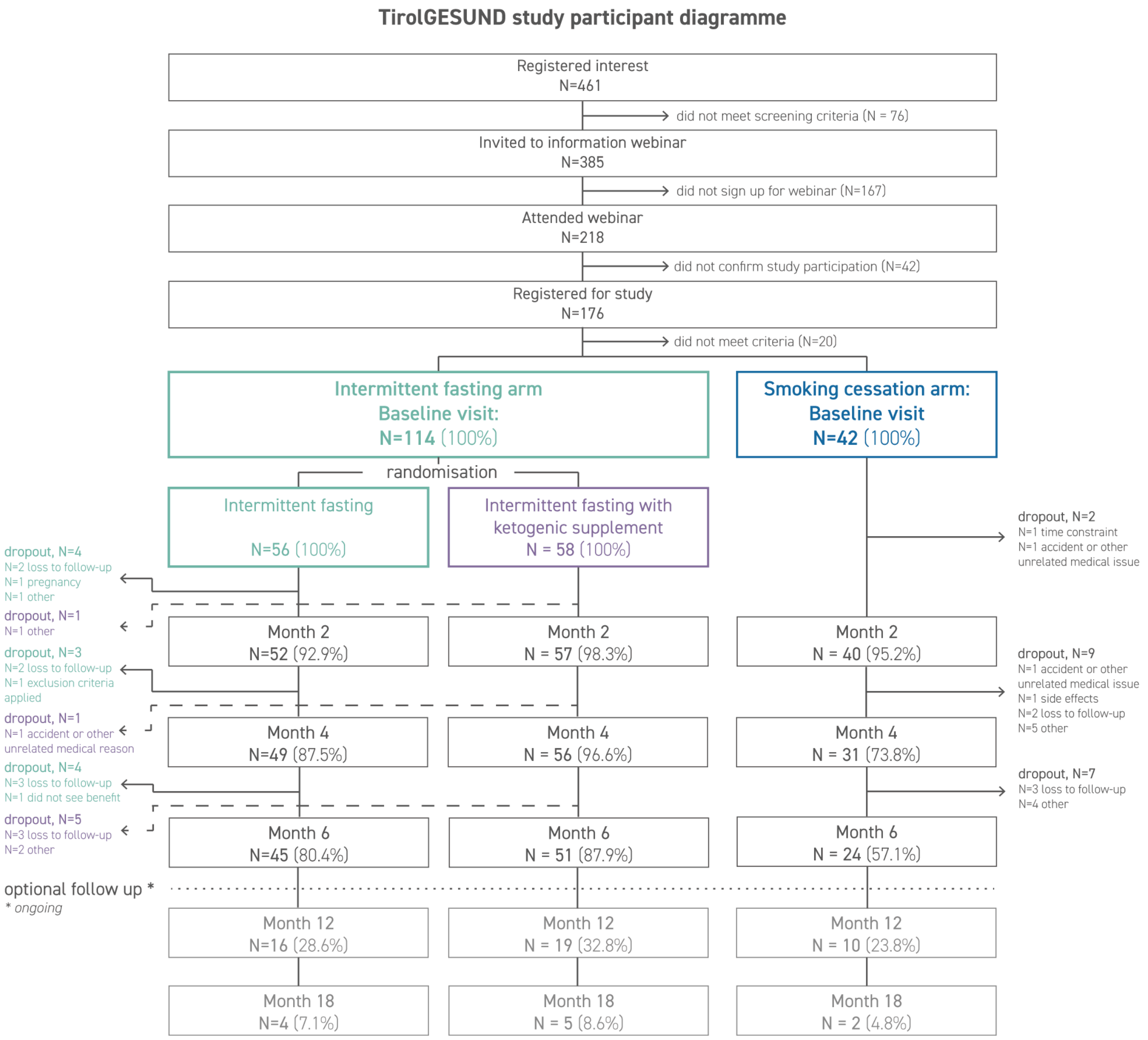


**Figure S1. Overview of the TirolGESUND participant flow.**

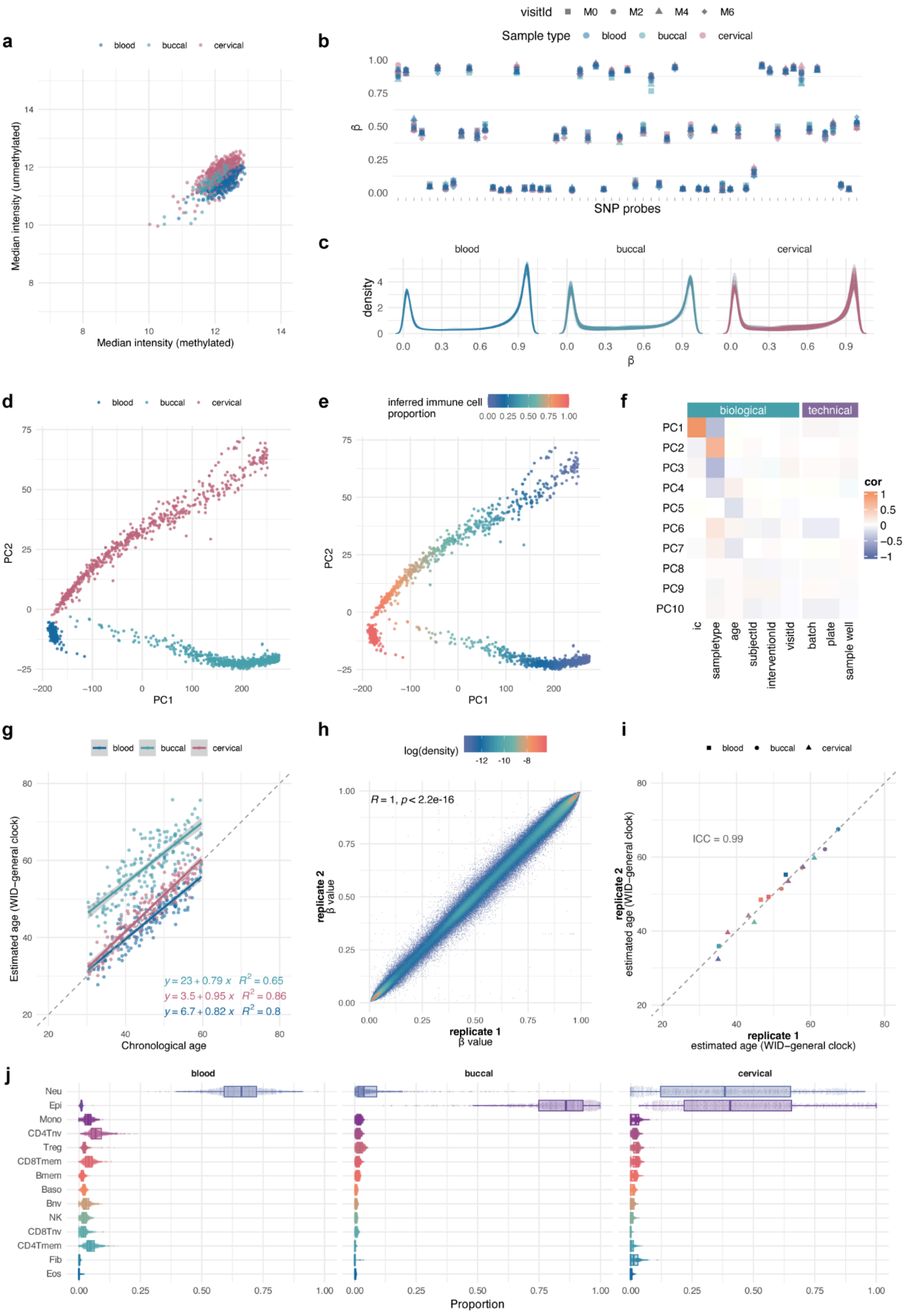

**Figure S2. Technical validation of methylation data.**

**a** Median intensity value for methylated and unmethylated samples (separate for blood, buccal, and cervical samples. **b** Example of single nucleotide polymorphism values across multiple sample types and timepoints used for quality control. **c** Beta methylation value density plots for all samples. d Principal component analysis shows distinct separation of principal components 1 and 2 (PC1, PC2) by sample type. **e** Inferred immune cell proportion of samples is strongly associated with principal component 1 (PC1). **f** Correlation of principal components with biological and technical factors. Samples from the same individual and sample type were processed in the same batch and plate. **g** Estimated age based on the WID general clock reveals good correlation with chronological age. As previously observed, age estimations using the WID-general clock are less accurate in buccal samples. Correlation value indicates Pearson's r. **h** Correlation of beta methylation values in a technical replicate (representative cervical sample). **i** Correlation of estimated age (WID-general clock) reveals high precision and repeatability in technical replicates. **j** Estimated cell type proportions using hierarchical EpiDISH (hierarchical epigenetic dissection of intrasample heterogeneity) with the centEpiFibIC.m and cent12CT.m reference matrices.

**Abbreviations:** PC, principal component. Neu, neutrophils. Epi, epithelial cells. Mono, monocytes. CD4Tnv, naive $CD4^+$ T cells. Treg, regulatory T cells. CD8Tmem, memory $CD8^+$ T cells. Bmem, memory B cells. Baso, basophils. Bnv, naive B cells. NK, NK T cells. CD8Tnv, naive $CD8^+$ T cells. CD4Tmem, memory $CD4^+$ T cells. Fib, fibroblasts. Eos, eosinophils.

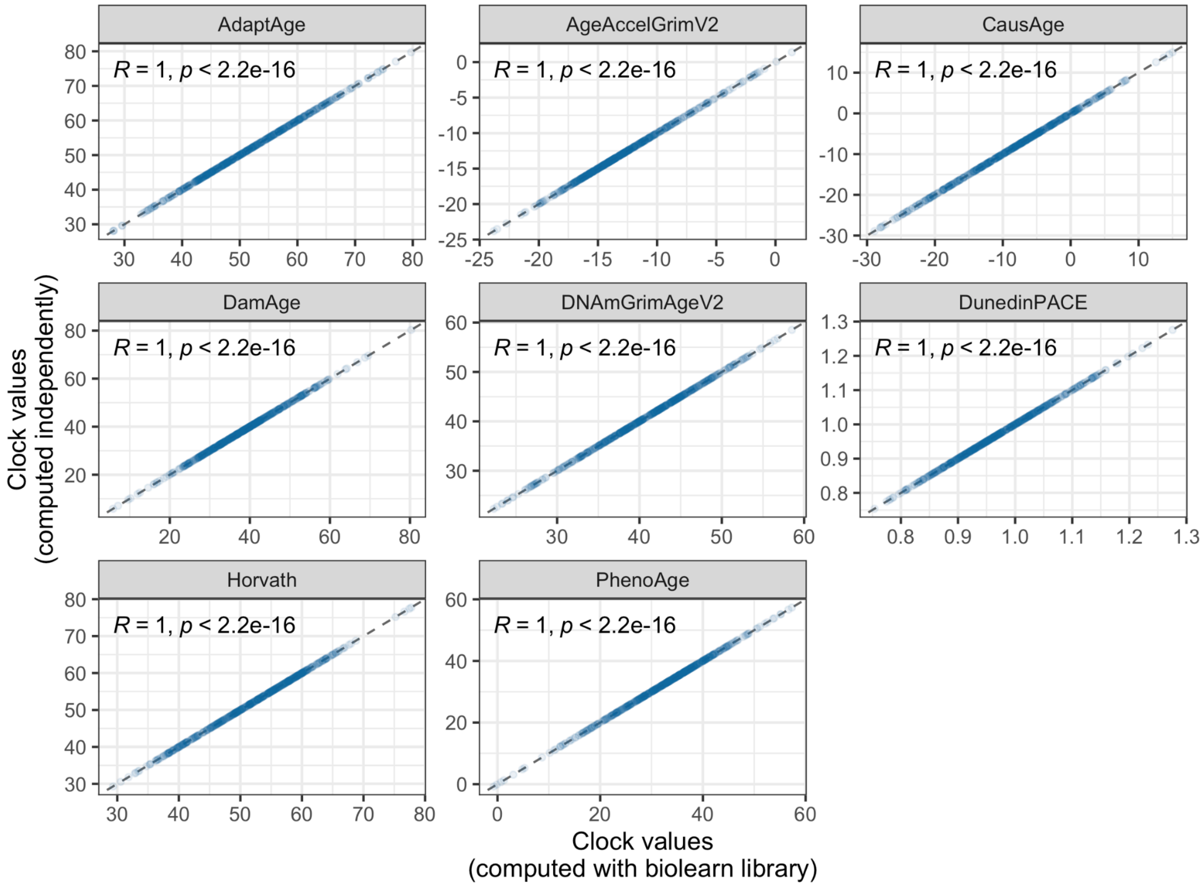


**Figure S3. Technical validation of external biomarker value algorithms.**
Biomarkers were computed independently using coefficients and algorithms from supplementary materials of respective original publications. Biomarker validity was validated using their calculation with the biolearn python library.
Correlation coefficient indicated is the Pearson correlation.

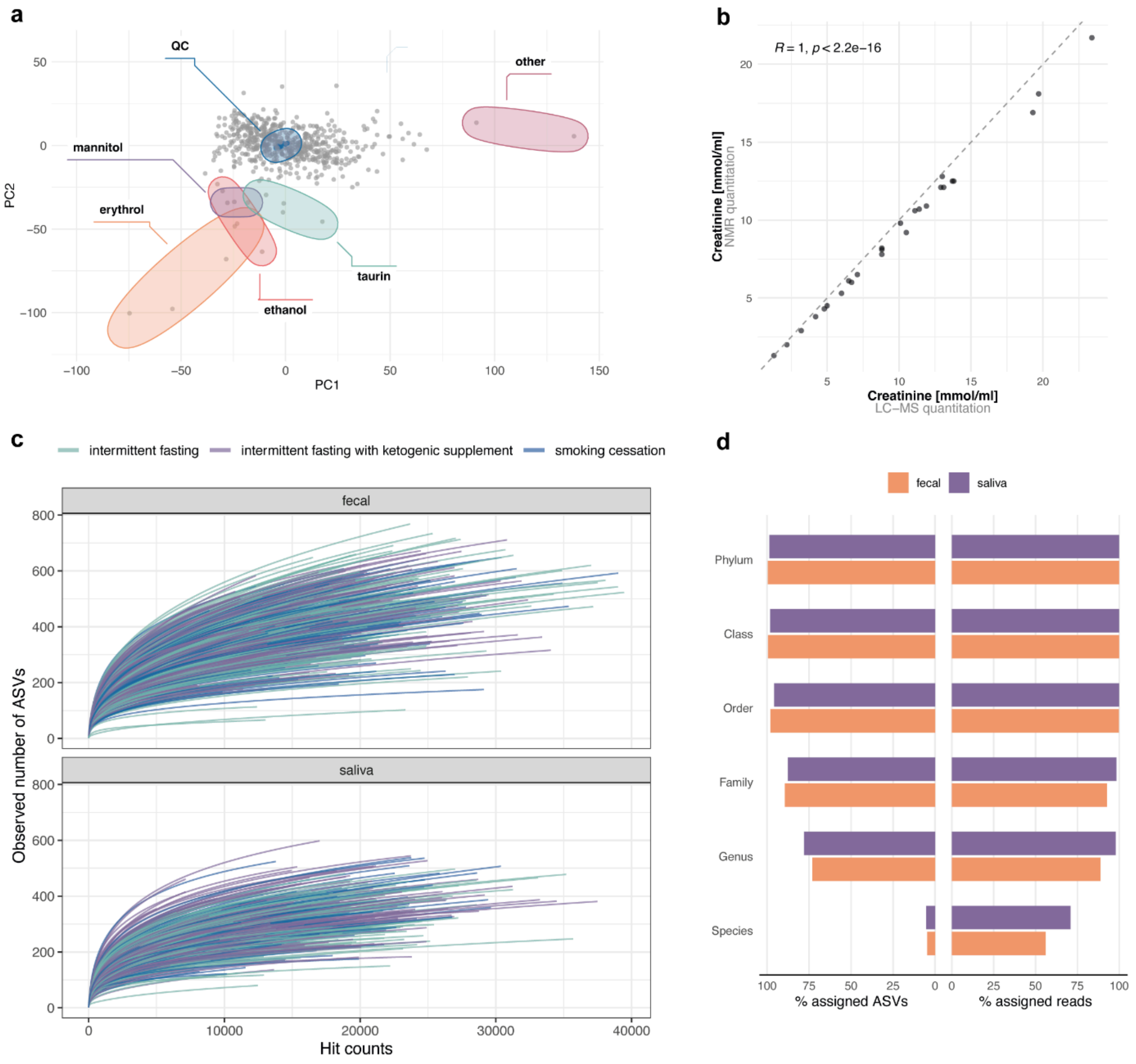


**Figure S4. Technical validation of metabolomics and microbiome data.**
**a** Principal component analysis of urine metabolome data reveals general clustering around quality control (QC) samples. Individual outliers have been identified and could be allocated to specific compounds, including sweeteners. **b** Correlation of quantitation by nuclear magnetic resonance and liquid chromatography-mass spectrometry. Correlation coefficient indicates Pearson's r. **c** Rarefaction curves for the prokaryotic amplicon sequence variants (ASVs) detected in the faecal and saliva microbiomes. Species richness estimated by the number of observed ASVs (Y-axis) is plotted against the number of subsampled read counts (X-axis) for each sample. Samples from the intermittent fasting and smoking cessation arms are depicted in blue and green, respectively. **d** Silva-based taxonomy assignment rate at different taxonomic levels for the amplicon sequence variants (ASVs) detected across the two study arms in the microbiome profiles. Top (purple) and bottom (orange) bars in each group depict the detection in saliva and faecal samples, respectively. Both percent of assigned unique reads (left-hand) and the respective non-unique counts (right-hand) are given, with non-prokaryotic ASVs and reads excluded from the total 100%.
**Abbreviations:** PC, principal component. NMR, nuclear magnetic resonance. LC-MS, liquid chromatography-mass spectrometry. ASV, amplicon sequence variant.

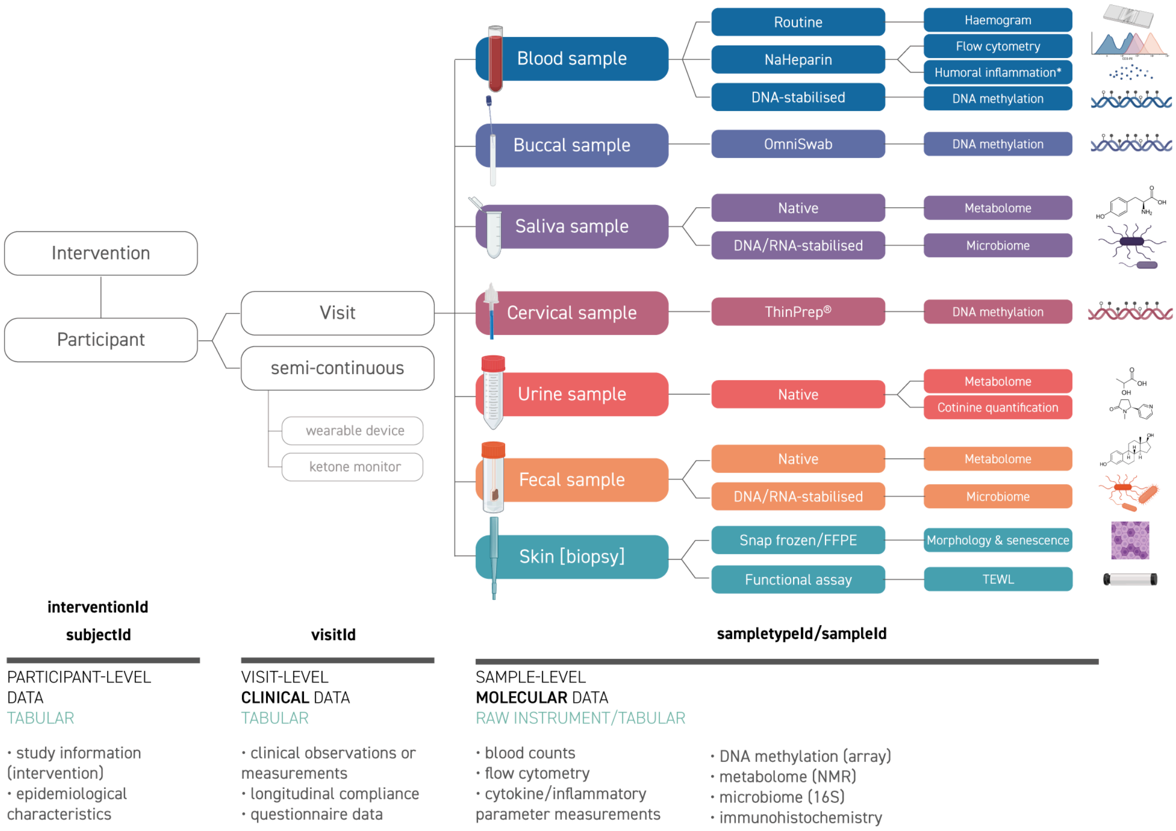


**Figure S5. Data structure overview.**
Data consist of multiple levels, including non-time-dependent baseline data such as epidemiological characteristics (tabular), time-dependent phenotypic information (including clinical) (tabular), or time-dependent sample-level molecular data (tabular or raw instrument data). Data are linked via distinct identifiers, mainly interventionId, subjectId, visitId, sampletypeId, and sampleId: a participant is identified by the subjectId, consisting of the interventionId (single letter I, intermittent fasting, K, intermittent fasting and ketogenic supplement, or S, smoking cessation) and a three-digit number. Visits are denoted by a visitId (M0, M2, M4, M6). Samples types are denoted by sampletypeIds. Samples and data are linked via subjectId (baseline or participant-level phenotypic data), subjectId and visitId, (visit-level phenotypic or clinical data), or sampleId (concatenating subjectId, visitId, and sampletypeId).
**Abbreviations:** TEWL, transepidermal water loss assay.

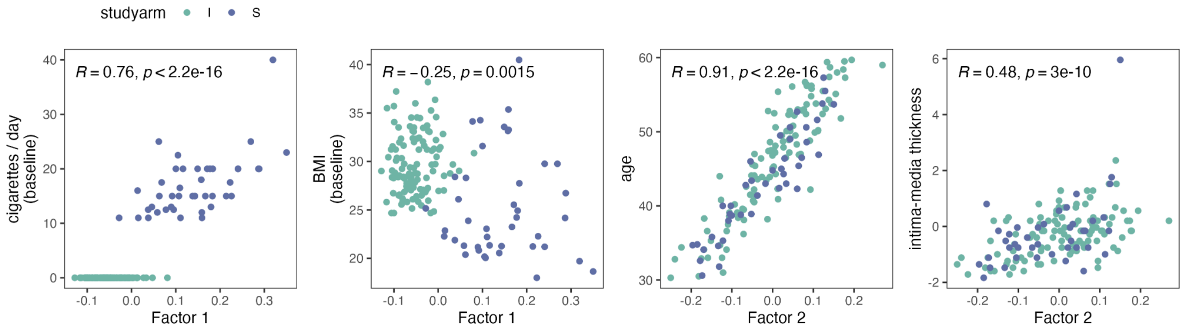


**Figure S6. Association of latent factors 1 and 2 deduced from a MOFA analysis at baseline with smoking, body-mass index, age, and intima-media thickness.**

**a** Correlation of Factor 1 with cigarettes per day at baseline. **b** Correlation of Factor 1 with body-mass index (BMI) at baseline. **c** Correlation of Factor 2 with age at baseline. **d** Correlation of Factor 2 with intima-media thickness at baseline.

Correlation coefficients indicate Spearman's rho.

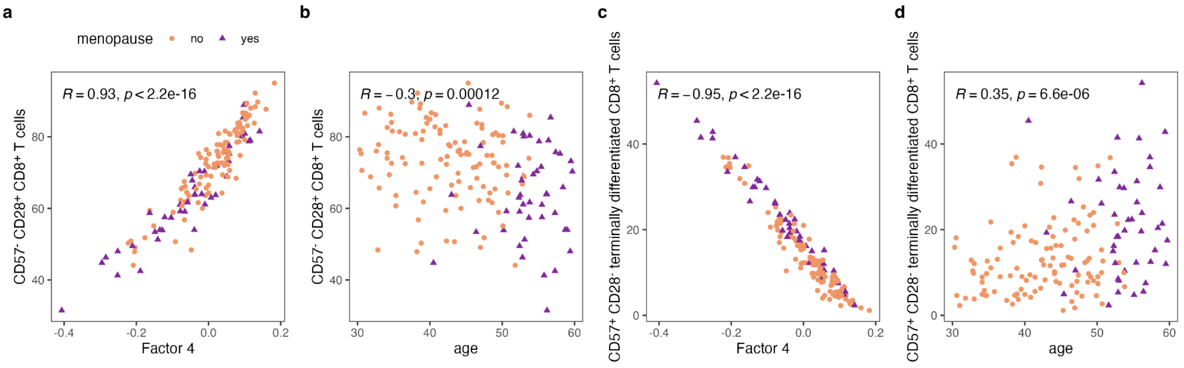


**Figure S7. Baseline MOFA reveals menopause-associated immune remodelling.**

**a** Correlation of CD57$^-$ CD28$^+$ CD8$^+$ T cells with factor 4 by menopausal status.

**b** Correlation of CD57$^-$ CD28$^+$ CD8$^+$ T cells with age by menopausal status.

**c** Correlation of CD57$^+$ CD28$^-$ terminally differentiated CD8$^+$ T cells with factor 4 by menopausal status.

**d** Correlation of CD57$^+$ CD28$^-$ terminally differentiated CD8$^+$ T cells with age by menopausal status.

Correlation coefficients indicate Spearman's rho.

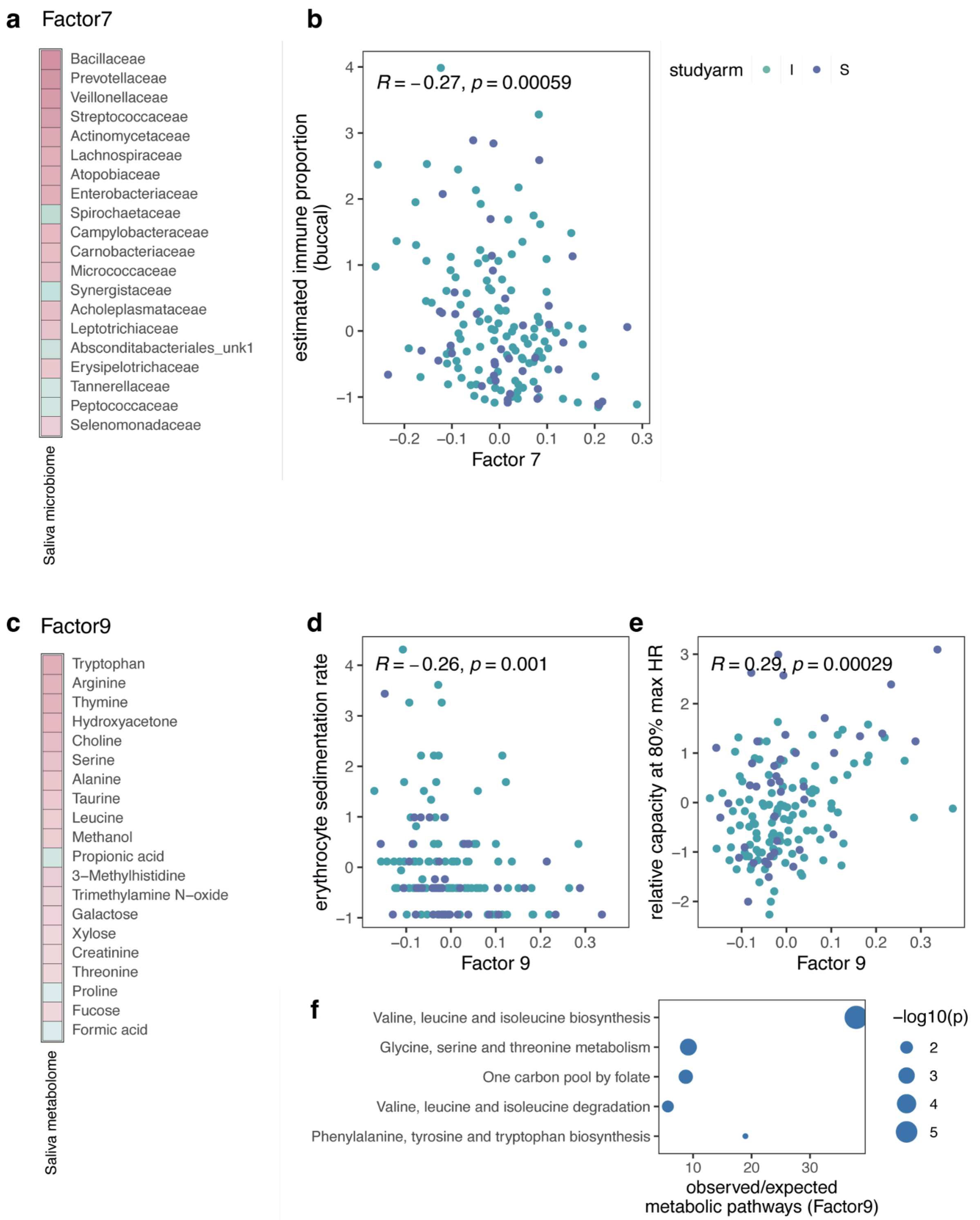


**Figure S8. Oral microbiota at baseline are associated with immune infiltration while oral metabolites associate with systemic inflammation and exercise.**

**a** Top weights for Factor 7 in baseline MOFA. **b** Correlation of Factor 7 with scaled and centred immune cell proportion in buccal samples, estimated by methylation. **c** Top weights for Factor 9 in baseline MOFA. **d** Association of Factor 9 with erythrocyte sedimentation rate. **e** Association of Factor 9 with relative capacity at 80% of the maximum heart rate. **f** Enrichment of top saliva metabolites in Factor 9 (|weight| ≥0.35).

Correlation coefficients indicate Spearman's rho.

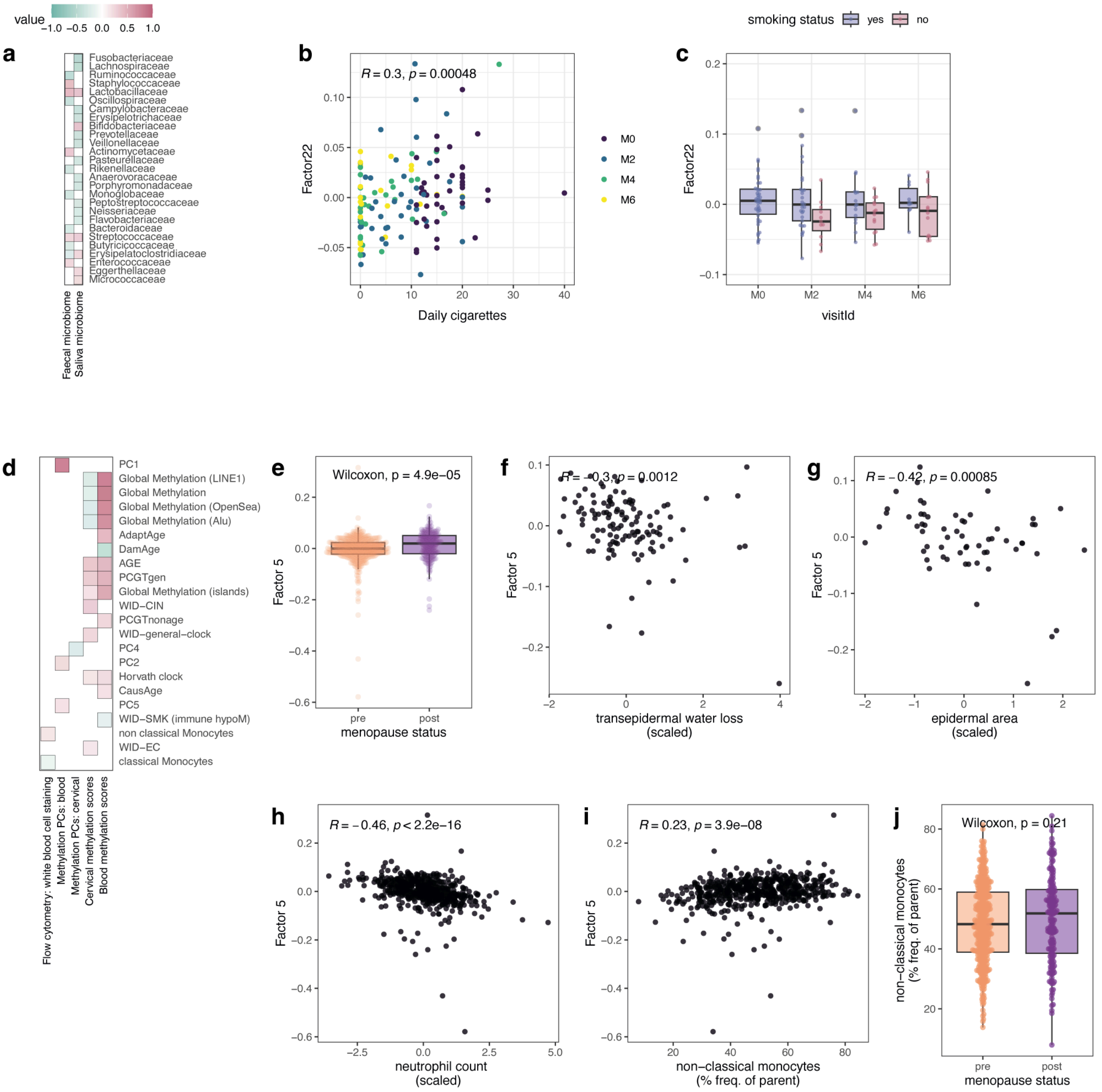


**Figure S9. Microbiome and immune features associated with smoking, menopause, and skin barrier.**

**a** Top features contributing to Factor 22. **b** Factor 22 correlates with daily cigarette consumption. **c** Factor 22 decreases after stopping smoking in the smoking cessation arm.

**d** Top features contributing to Factor 5. The scale is the same as in a. **e** Factor 5 is significantly increased in postmenopausal women. Factor 5 negatively correlates with transepidermal water loss (**f**), epidermal area (**g**), and neutrophils (**h**) and positively correlates with non-classical monocytes (**i**). **j** Non-classical monocytes are elevated after menopause, but this does not reach significance.

Correlation coefficients in b and d-i are Spearman correlation coefficients.

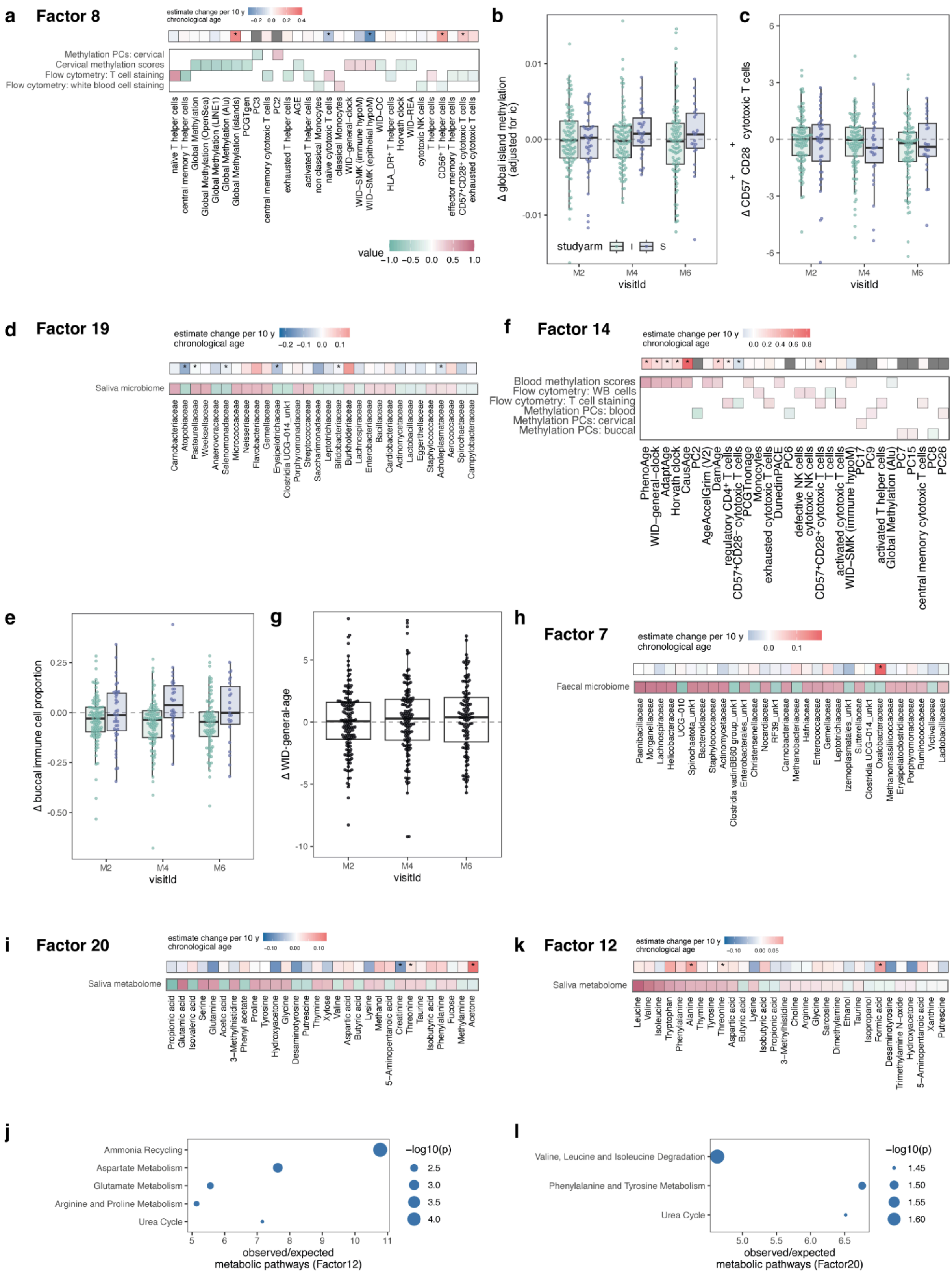


**Figure S10. Global changes in immune cells, ageing features, faecal microbiome, and oral metabolome.**

**a** Top weights for Factor 8 in longitudinal MEFISTO analysis and their baseline association with chronological age in a multi-omic-wide association study (MOWAS; excluding methylation PCs). **b** Change in cervical global island methylation from baseline by study group. **c** Change in $CD57^{+}CD28^{+}$ cytotoxic T cells from baseline by study group. **d** Top

weights for Factor 19 in longitudinal MEFISTO and their baseline association with chronological age in the MOWAS. **e** Change in buccal immune cell composition from baseline by study group. **f** Top weights for Factor 14 in longitudinal MEFISTO and their baseline association with chronological age in the MOWAS. **g** Change in WID general age from baseline shows gradual increases in epigenetic age. **h** Top weights for Factor 7 in longitudinal MEFISTO and their baseline association with chronological age in the MOWAS. **i** Top weights for Factor 20 in longitudinal MEFISTO and their baseline association with chronological age in the MOWAS. **j** Enrichment analysis for top weights in Factor 20 (|cor|>35). **k** Top weights for Factor 7 in longitudinal MEFISTO and their baseline association with chronological age in the MOWAS. **l** Enrichment analysis for top weights in Factor 12 (|cor|>35).
Colour scales for factor weights are consistent between a, d, f, h, i, and k.
Baseline associations with chronological age were assessed using linear models (see MOWAS section). Unadjusted p values < 0.05 are denoted with *.

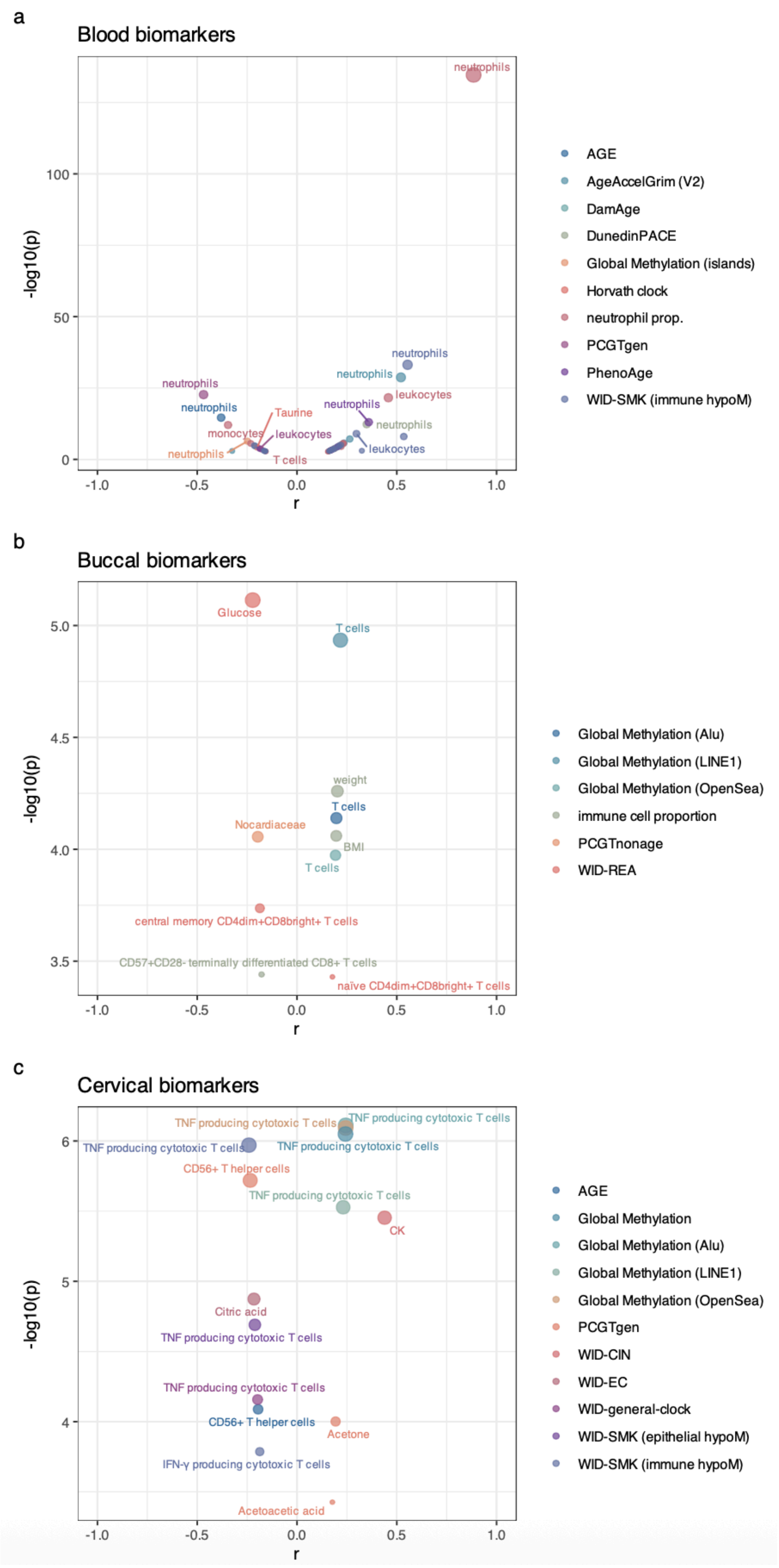


**Figure S11. Repeated measures correlation overview.**

Volcano plots of **a** blood biomarker, **b** buccal biomarker, and **c** cervical biomarker associations at fdr<0.05.

Correlation coefficients indicate repeated measures correlation ($r_{rm}$), p values indicate unadjusted p values.
